# Decomposition of retinal ganglion cell electrical images for cell type and functional inference

**DOI:** 10.1101/2023.11.06.565889

**Authors:** Eric G. Wu, Andra M. Rudzite, Martin O. Bohlen, Peter H. Li, Alexandra Kling, Sam Cooler, Colleen Rhoades, Nora Brackbill, Alex R. Gogliettino, Nishal P. Shah, Sasidhar S. Madugula, Alexander Sher, Alan M. Litke, Greg D. Field, E.J. Chichilnisky

## Abstract

**Objective:** Identifying neuronal cell types and their biophysical properties based on their extracellular electrical features is a major challenge for experimental neuroscience and for the development of high-resolution brain-machine interfaces. One example is identification of retinal ganglion cell (RGC) types and their visual response properties, which is fundamental for developing future electronic implants that can restore vision.

**Approach:** The electrical image (EI) of a RGC, or the mean spatio-temporal voltage footprint of its recorded spikes on a high-density electrode array, contains substantial information about its anatomical, morphological, and functional properties. However, the analysis of these properties is complex because of the high-dimensional nature of the EI. We present a novel optimization-based algorithm to decompose electrical image into a low-dimensional, biophysically-based representation: the temporally-shifted superposition of three learned basis waveforms corresponding to spike waveforms produced in the somatic, dendritic and axonal cellular compartments.

**Results:** The decomposition was evaluated using large-scale multi-electrode recordings from the macaque retina. The decomposition accurately localized the somatic and dendritic compartments of the cell. The imputed dendritic fields of RGCs correctly predicted the location and shape of their visual receptive fields. The inferred waveform amplitudes and shapes accurately identified the four major primate RGC types (ON and OFF midget and parasol cells) substantially more accurately than previous approaches.

**Significance:** These findings contribute to more accurate inference of RGC types and their original light responses based purely on their electrical features, with potential implications for vision restoration technology.

## Introduction

The increasing scale and density of multi-electrode neural recordings opens up the possibility of using *electrical imaging* – spatiotemporal analysis of extracellularly recorded voltage waveforms produced by a cell – for diverse applications in neuroscience and neuroengineering. An important example is the development of epiretinal implants, which are designed to restore vision by evoking spikes in retinal ganglion cells (RGCs) in the degenerated retina, thereby conveying artificial visual signals to the brain. Because the ∼20 RGC types in the human and non-human primate retina have very different light response properties [Roska 2014], targeting the distinct RGC types independently is fundamental to accurately replicating the natural neural code of the retina. This requires an implant that can use electrical recordings in a blind retina to identify the distinct RGC types. The electrical image (EI) of a RGC – the mean spatiotemporal voltage footprint of its recorded spikes on a high-density multi-electrode array (MEA) – contains substantial information about the morphology, location, and biophysical properties of the cell that could potentially be used to infer its cell type and receptive field. However, because EIs are high-dimensional and complex, using them for these purposes has been difficult [Richard 2015, Zaidi 2023]. Thus, a low-dimensional and interpretable representation of the EI could be useful for probing the cell type and function of RGCs for future vision restoration efforts, and could also be valuable for electrical images recorded in various parts of the brain.

Here, we present an optimization-based approach to decompose the EIs of RGCs into sums of temporally-shifted learned somatic, dendritic, and axonal basis waveforms. The decomposition is low-dimensional and interpretable, consisting of the learned cellular compartment basis waveform shapes and their respective amplitudes and time shifts for each recording electrode on the MEA. We first validate the algorithm by comparing the spatial arrangement of the EI decomposition with labeled confocal micrographs of the recorded cell to demonstrate correspondence with its geometric properties. We then show that RGC receptive field centers and shapes can be inferred from the dendritic component of the decomposition. Finally, we demonstrate that the fitted cellular compartment waveforms and amplitudes each systematically vary with RGC functional type, and that these properties alone can be used to accurately identify the four numerically-dominant primate RGC types. Together, these results demonstrate the use of the EI decomposition to systematically characterize the anatomical and physiological properties of RGCs, and thus could improve the functionality and fidelity of retinal implants for vision restoration and provide a framework for analysis of electrical images in other parts of the nervous system.

## Results

### Decomposition of electrical images into somatic, dendritic and axonal components

We exploited the features of electrical images (EIs) of primate retinal ganglion cells (RGCs) recorded on a multi-electrode array (MEA) to reveal biophysical properties of neurons, by developing a decomposition based on cellular compartments. The voltage waveforms recorded extracellularly arise from time-varying charge sources and sinks in extracellular medium induced by trans-membrane currents during action potential propagation. The EI, or average spatiotemporal voltage recorded during a spike, provides a view of the spatial and temporal structure of these aggregated biophysical quantities (Fig. 1A-B). The shapes of somatic, dendritic, and axonal waveforms in the EI are stereotyped [Petrusca 2007], with somatic waveforms having biphasic shape with a strong early negative peak (e.g. waveform 1 in Fig. 1B), dendritic waveforms having a biphasic shape with a strong early positive peak (e.g. waveform 2 in Fig. 1b), and axonal waveforms having a strong negative peak and overall triphasic shape indicative of a traveling wave (e.g. waveforms 5 and 6 in Fig. 1B) [Henze 2000, Gold 2006, Gold 2009]. However, many of the voltage waveforms comprising the EI consist of superpositions of signals from multiple parts of the cell (e.g. waveforms 3 and 4 in Fig. 1B), due to the morphology of RGCs, the spatial blurring inherent in extracellularly-recorded signals, and the spatiotemporally propagating nature of action potentials. This creates a large diversity of waveform shapes and complicates analysis of spatiotemporal structure in the EI.

**Figure 1.**
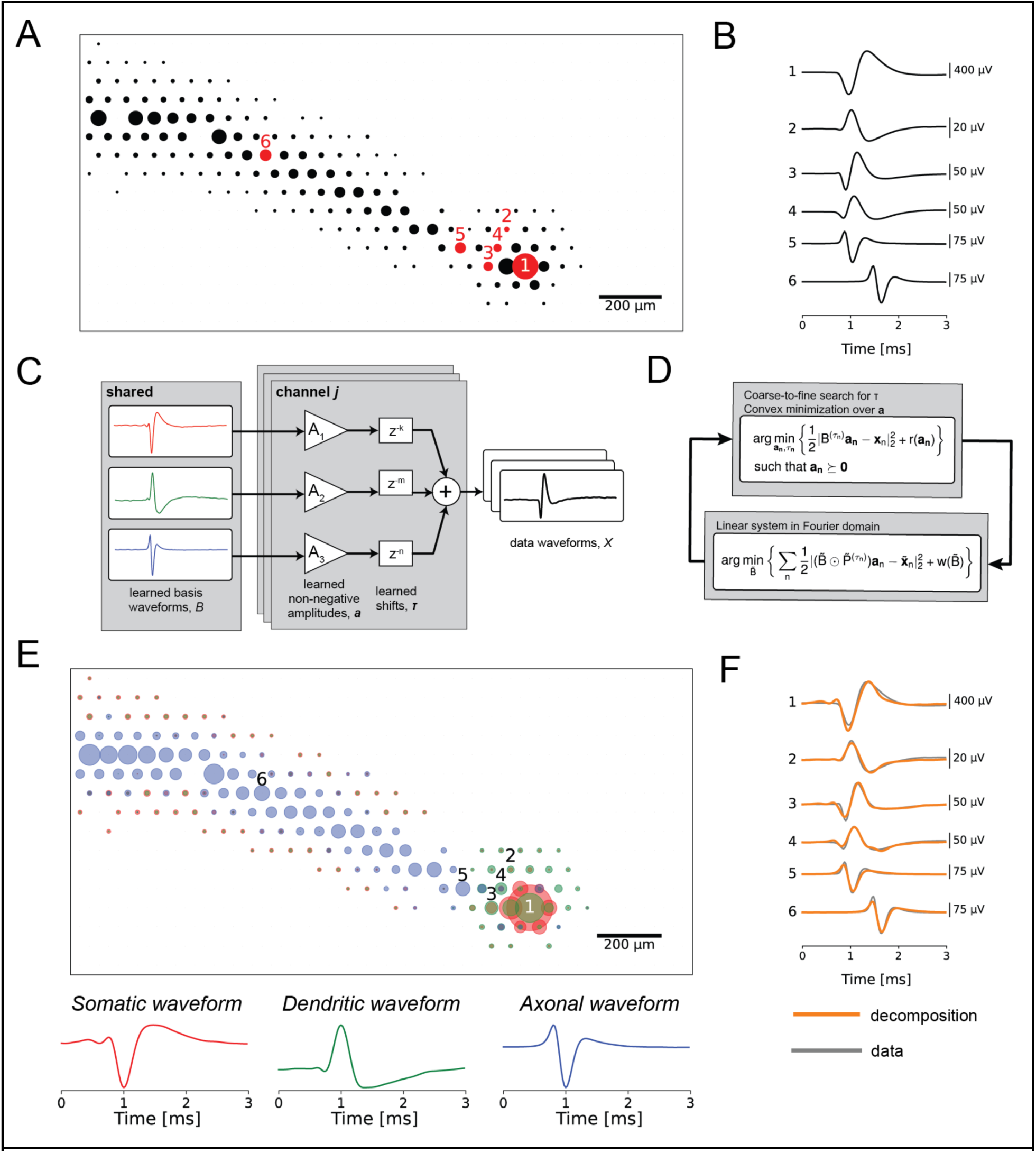
**(a)** Spatial component of the electrical image (EI) for an ON parasol cell. The diameters of the circles correspond to the amplitude of the signal recorded on each recording electrode of the MEA. **(b)** Extracellularly-recorded waveforms from the EI. The numerical labels correspond to the electrodes numbered in (a). Waveform 1 is a stereotypical somatic waveform, which is biphasic and consists of a strong fast negative peak followed by a slower positive peak. Waveform 2 is a stereotypical dendritic waveform, consisting of a strong fast positive peak followed by a slower negative peak. Waveforms 5 and 6 are stereotypical axonal waveforms, consisting of a strong negative peak and an overall triphasic shape, indicative of a traveling wave. Waveforms 3 and 4 are examples of superpositions, as their shapes do not match any of the stereotyped basis waveforms. **(c)** Signal model for the EI decomposition algorithm. The recorded data waveform on each electrode is modeled as the non-negatively weighted sum of temporally-shifted compartment basis waveforms. Distinct amplitudes and shifts are learned for each recording electrode, while the basis waveforms are shared across all electrodes for a given cell. **(d)** Iterative optimization procedure for fitting the EI decomposition. The algorithm alternates between an amplitude and shift fitting step, and a waveform shape optimization step. The variable names are matched with panel (c), and full descriptions of the mathematical approach and optimization procedure are provided in the Methods. **(e)** Decomposition of the ON parasol cell from (a). Top: spatial representation of the learned amplitudes of the EI decomposition. The color of each circle corresponds to the compartments of the decomposition (red for soma, green for dendrite, blue for axon), and the diameters of the circles correspond to the fitted amplitudes. Note that the spatial spread of the signal recorded by the MEA substantially exceeds the physical size of the cell, a well established property of MEA retinal recordings. Bottom: fitted basis waveforms for the soma, dendrites, and axon, respectively. **(f)** Overlay of the waveforms reconstructed from the decomposition (orange) on top of the original data waveforms (gray). The decomposition accurately represented the shape diversity in the data waveforms using only three learned basis waveforms.

To distinguish the underlying cellular compartments, the EI decomposition represents the recorded extracellular voltage waveform of a cell on each electrode as the shifted and scaled superposition sum of three learned basis waveforms, putatively corresponding to dendritic, somatic, and axonal compartments (Fig. 1C). These learned basis waveforms are shared among all recording electrodes, but are fitted separately for every cell, allowing the decomposition to capture physiological differences between cells and cell types.

The EI decomposition is fitted by minimizing the mean square error between the recorded EI waveforms and the waveforms reconstructed from the decomposition. The learned weights on the basis waveforms are forced to be non-negative, as they represent contributions of physical compartments of the cell to the recorded electrical signal. The overall problem is non-convex, and is solved approximately using a novel algorithm for shifted semi-nonnegative matrix factorization (see Methods), related to nonnegative matrix factorization [Lee 2000] and shifted nonnegative matrix factorization [Morup 2007]. Specifically, the decomposition algorithm iteratively alternates between two steps: (1) an amplitude and time shift fitting step, holding the basis waveforms fixed; and (2) a basis waveform fitting step, holding the amplitudes and time shifts fixed. The decomposition fitting objective includes additional regularization terms to induce sparsity in the fitted amplitudes and to constrain the shapes of the learned basis waveforms. Details of the regularization and selection of associated hyperparameters are provided in the Methods.

The decomposition accurately represented the diverse waveforms in the EI. The EI was decomposed into a ball corresponding to cell soma, a region surrounding the soma representing the dendritic field, and a linearly propagating signal representing the axon (Fig. 1E, top). The shapes of the learned basis waveforms (Fig. 1E, bottom) approximately matched the waveform shapes expected for somatic, dendritic, and axonal compartments based on first principles [Petrusca 2007]. Finally, the data waveforms (gray lines in Fig. 1F) were accurately captured by shifted superpositions of the three compartment basis waveforms (orange lines in Fig. 1F), even though their shapes differed markedly from the individual basis waveforms, demonstrating the representational power of the decomposition.

### Decomposition components approximately match cellular morphology

To evaluate whether the components of the fitted decomposition corresponded to the morphological features of RGCs, the correspondence between the decomposition and fluorescently labeled cells was tested. The decomposition somatic component was validated by comparison with manually-labeled ON and OFF parasol cell bodies visualized in confocal micrographs (Fig. 2A). Decomposition somatic centers were calculated as the signal-weighted position of the largest-amplitude somatic electrode and its six nearest neighbor electrode [Zaidi 2023], while ground truth cell body locations in the micrographs were identified by staining with a dye labeling HCN1, a channel protein thought to be selectively expressed in the parasol cell body [Goloshchapov 2009, Li 2015]. The staining procedure (see Methods) required removing the retina from the MEA prior to imaging, potentially warping the geometry of the retina in the images relative to the electrophysiological recording. Therefore, an approximate geometric alignment between the coordinates of the recording electrodes of the MEA and micrograph coordinate space was estimated by aligning vasculature and other large identifiable features (see Methods). The decomposition algorithm effectively identified somatic centers from the EI, with median errors of 29.6 μm and 28.0 μm for the ON parasol cells and OFF parasol cells respectively, comparable to the 30 μm spacing between recording electrodes (Fig. 2B). Due to the strong likelihood of mechanical warping of the retina, these errors can be interpreted as a lower bound on the precision of the method. Note that this analysis did not substantially improve upon simpler *ad hoc* methods for identifying the soma center from the EI [Li 2015]; thus, this result serves as a validation rather than a novel finding.

**Figure 2.**
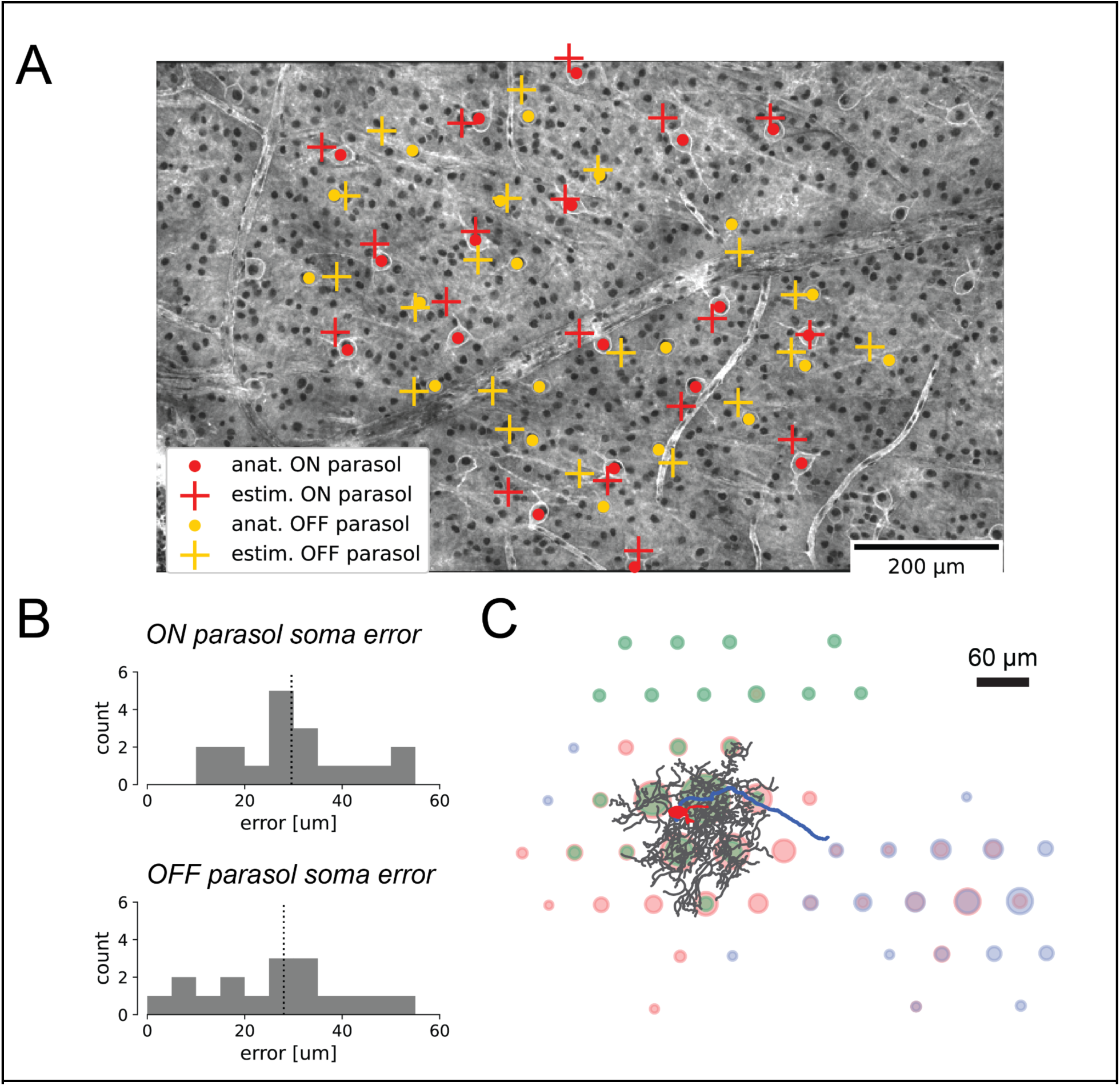
Alignment of the decomposition somatic and dendritic components with labeled micrographs of the retina. **(A)** Alignment of decomposition-estimated soma centers with anatomically-labeled soma centers for ON and OFF parasol cells (see legend). **(B)** Histograms quantifying soma center localization error for ON and OFF parasol RGCs. The median localization error was 29.6 um for ON parasols and 28.0 um for OFF parasols, comparable to the 30 um recording pitch of the MEA. **(C)** EI decomposition of a single rat RGC, where the color of each circle at each electrode corresponds to the dominant compartment of the decomposition at that electrode (red for soma, green for dendrite, blue for axon), and the diameters of the circles correspond to the recorded spike amplitudes. The decomposition is overlaid with the traced cell body (red), dendritic tree (black), and axon (blue) characterized by staining and imaging the retina.

The correspondence of the somatic and dendritic components of the decomposition was further evaluated using projection targeting with photo-tagging. This approach enables physiological identification of a cell using optogenetically-derived responses and provides an extraordinarily detailed view of the cell’s morphological features using fluorescence imaging simultaneously with recording, enabling precise geometric alignment between the photomicrographs and the recorded electrical signals. Using this technique, a single rat RGC was imaged, and the cell body, dendritic tree, and axons were traced and overlaid onto the fitted EI decomposition (Fig. 2C). Comparison of the decomposition with the traced cellular compartments revealed a general correspondence between the decomposition and the anatomical layout of the cell. Notably, however, the spatial extents of the decomposition-estimated components for each compartment were substantially larger than the traced cell geometries, suggestive of spatial blurring of the voltage signal captured by extracellular recording.

### Inferred dendritic fields strongly correlate with measured receptive fields

Precise knowledge of the receptive field location and shape for each RGC is needed for an epiretinal implant to effectively mimic the neural code [Shah 2020, Zaidi 2023]. Although these properties can be easily characterized using light stimulation in a healthy retina, they must be inferred from electrically-recorded signals in a retina lacking light responses. Previous work performed this inference using the somatic signal alone. However, because RGCs receive visual input through their dendrites, and dendritic fields are not necessarily symmetric about the cell soma, inclusion of the recorded dendritic signals could improve estimates of location and shape of RGC receptive fields. Due to the challenge of estimating the dendritic contributions to EIs, however, this possibility has been difficult to evaluate. The decomposition algorithm solves precisely this problem.

Inclusion of the dendritic centers significantly improved the accuracy of inferred receptive field locations. This was demonstrated in 29 preparations by comparing the fit quality of affine transformations mapping somatic centers to receptive field centers with that of affine transformations that additionally included the decomposition-estimated dendritic field centers. The dendritic field centers were estimated as the weighted centers-of-mass of the decomposition dendritic amplitudes, while the somatic centers were estimated as the weighted centers-of-mass of the decomposition somatic amplitudes over the largest amplitude somatic electrode and its six nearest neighbor electrodes [Zaidi 2023]. Receptive field centers were computed from the stimulus spike-triggered average obtained with white noise visual stimulation (see Methods). Distinct mappings were fitted for each cell type, because the relationships between the somatic, dendritic, and receptive field centers could differ by cell type due to biological differences or tissue distortions. To account for differences in the eccentricities of the recordings and across cell types, receptive field estimation error was normalized by the mean receptive field diameter. For nearly every experimental preparation, inclusion of the decomposition-estimated dendritic center reduced the estimation error for receptive field centers (Fig. 3), with median error reduction across preparations of 0.0737, 0.0864, 0.315, and 0.181 receptive field diameters for the ON parasol cells, OFF parasol cells, ON midget cells, and OFF midget cells, respectively. Comparisons of the error relative to receptive field size revealed that the inferred receptive field centers were more accurate for parasol RGCs than for midget RGCs, and that inclusion of the dendritic center reduced the error more for midget RGCs than for parasol RGCs, aligning with previous observations that the dendritic fields of midget RGCs can be more offset from the soma than those of parasol RGCs [Watanabe 1989].

**Figure 3.**
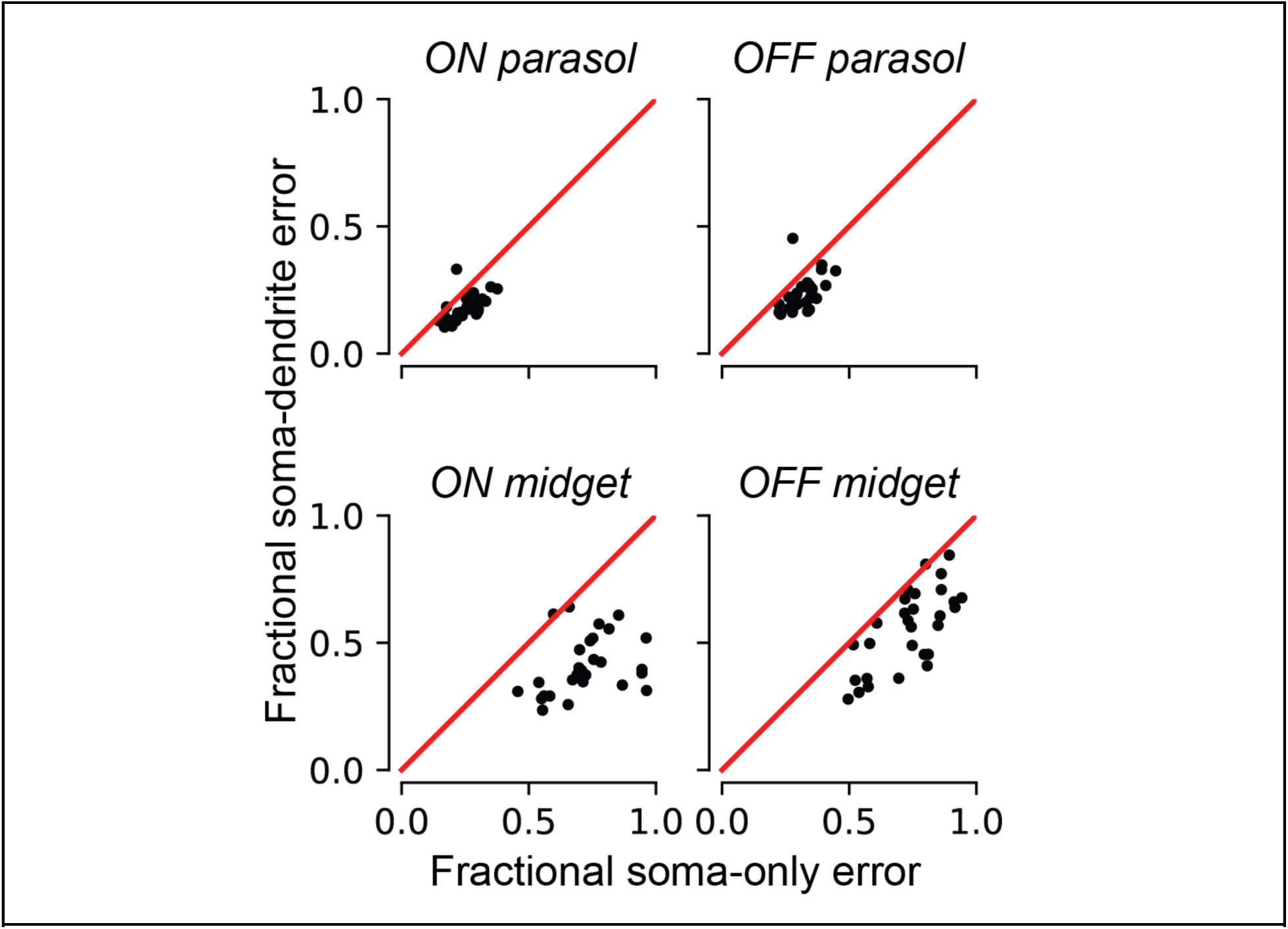
Comparison of inferred receptive field center using the location of the somatic center alone, and using both the somatic and dendritic field centers, for the four major cell types of the primate retina. Each point in the scatterplots corresponds to one experimental preparation. In total, 29 experimental preparations were analyzed. The y-axis describes the prediction error using both the somatic and dendritic field center, and the x-axis describes the prediction error using the somatic center alone. The magnitudes of the errors are expressed as a fraction of the mean receptive field diameter to facilitate comparison between preparations and cell types.

The dependence of the receptive field center estimates on the EI signal-to-noise ratio (SNR) was evaluated in one experimental preparation by artificially subsampling recorded spikes. EIs were recomputed using 100, 1000, and 10000 recorded spikes, corresponding to effective recording lengths of ∼10, ∼100, and ∼1000 seconds, respectively (Supplemental Figure S3B). Decompositions were refitted and the receptive field center estimation analysis was repeated separately for each cell type at each spike count level (Supplemental Figure S4). With the exception of ON midget RGC decompositions using 100 recorded spikes, the estimation error remained relatively unchanged, suggesting that receptive field estimation was robust to the amount of recorded data (and thus SNR) over a reasonable range.

To test whether the shapes of estimated RGC dendritic fields were informative about receptive field shape, the shapes of maximum-tiling dendritic field contours were compared with their measured receptive field counterparts. This comparison was performed using two large, lower-density RGC types, the OFF smooth monostratified cells [Rhoades 2019] and the putative broad thorny cells [Kling 2023], chosen because complete populations could be recorded, and because their large dendritic fields could easily be probed using a MEA with 60 μm recording electrode pitch. Maximum-tiling dendritic field contours were computed by blurring the decomposition dendritic fields for each cell into smooth 2D surfaces (see Methods), and then thresholding to maximize the total area covered by exactly one cell [Gauthier 2009]. The maximum-tiling dendritic field contours were then mapped from recording electrode coordinate space into stimulus space (see Methods), and overlaid on top of the maximum-tiling receptive field contours [Gauthier 2009] for comparison (Fig. 4A and 4C). Despite their irregular shapes, the dendritic and receptive field contours showed a striking degree of shape concordance in the two preparations tested, demonstrating a tight association between the structure of the RGC receptive field and the geometry of the dendritic field inferred by the decomposition.

**Figure 4.**
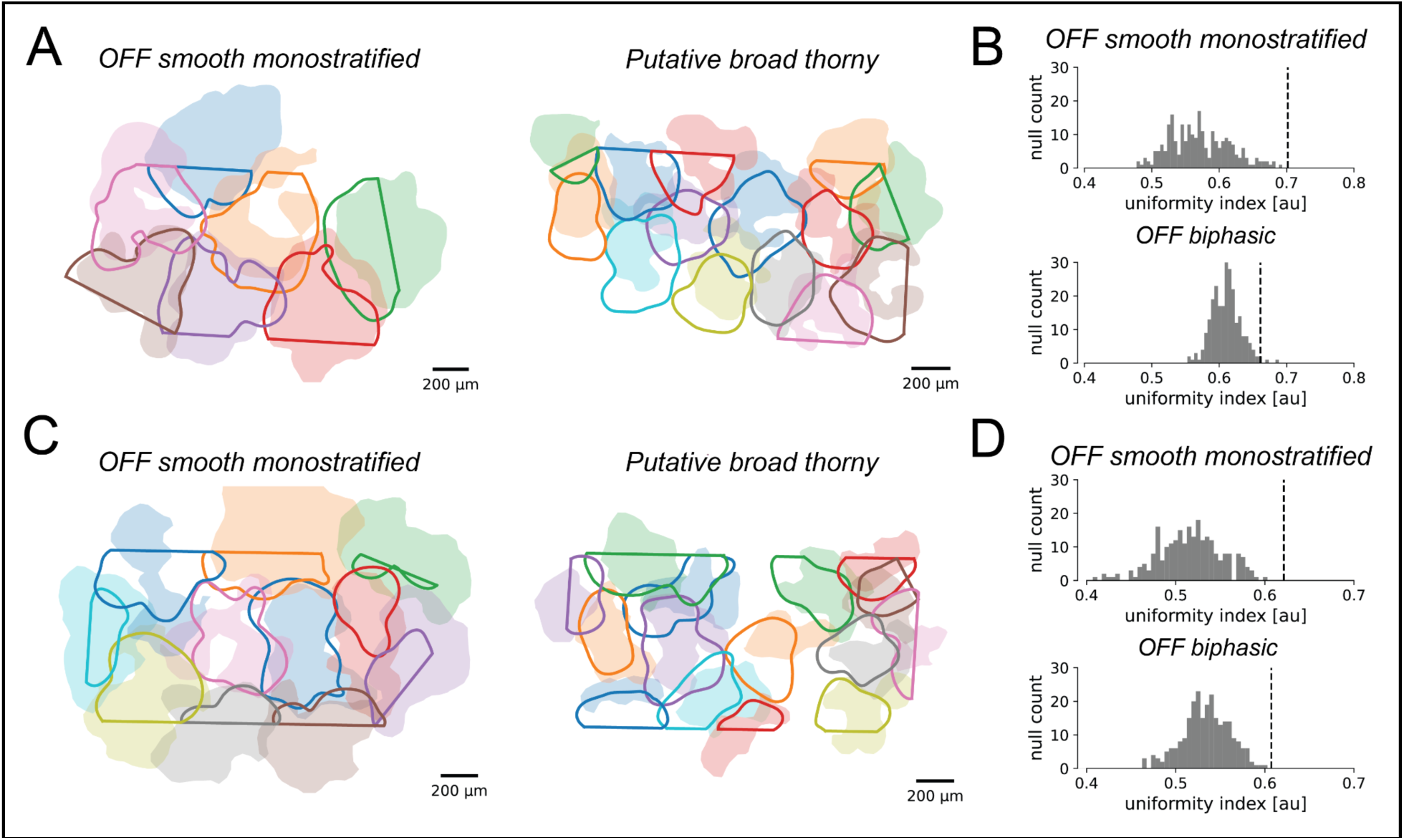
Dendritic field mosaics of OFF smooth monostratified and putative broad thorny ganglion cells, in two experimental preparations. **(A) and (C):** overlays of the dendritic field contours (solid lines) and the receptive field contours (shaded regions). The straight edges in the dendritic field contours are due to the boundaries of the MEA. **(B) and (D)**: Uniformity index (UI) characterizing the degree of dendritic field coordination, corresponding to the preparations in (A) and (C), respectively. The histograms show the null distribution of the UI (N=250), and the dotted lines mark the value of the data UI. P-values for mosaic tiling and coordination were < 1·10^-10^ and 0.008 for the OFF smooth monostratified cells and putative broad thorny cells respectively for the preparation in (a), and < 1·10^-10^ and 0.004 for the same respective cell types for the preparation in (b).

To further validate the relationship between RGC dendritic and receptive fields, the degree of coordination between dendritic fields of RGCs of the same type was evaluated to determine whether they shared the tiling and coordination properties characteristic of receptive fields [Gauthier 2009]. This was quantified using the uniformity index (UI), defined as the fraction of total MEA area contained within the dendritic contour of exactly one cell. The statistical significance of dendritic field coordination was evaluated by comparing the UI with a null distribution constructed by randomly rotating the dendritic fields of each RGC (see Methods). In each case, the degree of mosaic tiling and coordination in the observed data was highly significant (Fig. 4B and Fig. 4D). This is consistent with anatomical results demonstrating coordination of RGC dendritic fields [Wässle 1981, Dacey 1993]. However, the same analysis largely failed to recover statistically significant coordination in the inferred dendritic fields of ON and OFF parasol cells when recording at either 60 um electrode pitch or 30 um electrode pitch. This negative finding could be a consequence of the fact that adjacent parasol cell dendritic fields overlap significantly [Dacey and Brace 1992]. However, given that parasol receptive fields do exhibit interdigitation [Gauthier 2009], it is also potentially indicative of limitations in the spatial resolution of the decomposition algorithm and MEA recordings of RGC dendritic fields.

### Decomposition components distinguish different RGC types

Because the primate and human retina contains upwards of 20 functional cell types, each with distinct light response properties [Roska 2014], identification of the distinct RGC types in a degenerated retina without direct characterization of light responses may be important for the calibration of an epiretinal implant [Shah 2020]. To evaluate whether the decomposition can help to accomplish this identification, the separability of RGC functional types using the decomposition-fitted basis waveforms and amplitudes was examined.

The relationship between spike waveform shape and functional cell type was first characterized by applying principal components analysis (PCA) to the fitted basis waveforms of functionally-identified OFF parasol cells, OFF midget cells, OFF smooth monostratified cells [Rhoades 2019], and putative broad thorny cells [Kling 2023]. Cells from three experimental preparations from different animals were pooled together. To account for timing differences between preparations due to variability between animals and experimental conditions, the basis waveforms from different recordings were stretched in time to equalize their time scales for parasol cells across recordings and then temporally aligned (see Methods). PCA was performed separately for each compartment, and the waveforms were then projected onto the top two PCs (Fig. 5A). Across preparations, for each of the basis waveforms, cells of the same type tended to have similar waveforms, and cells of different types tended to have different waveforms, though substantial overlap remains. Thus, RGCs of different functional types can exhibit systematically different waveform shapes that are substantial enough revealed by a very simple analysis.

**Figure 5:**
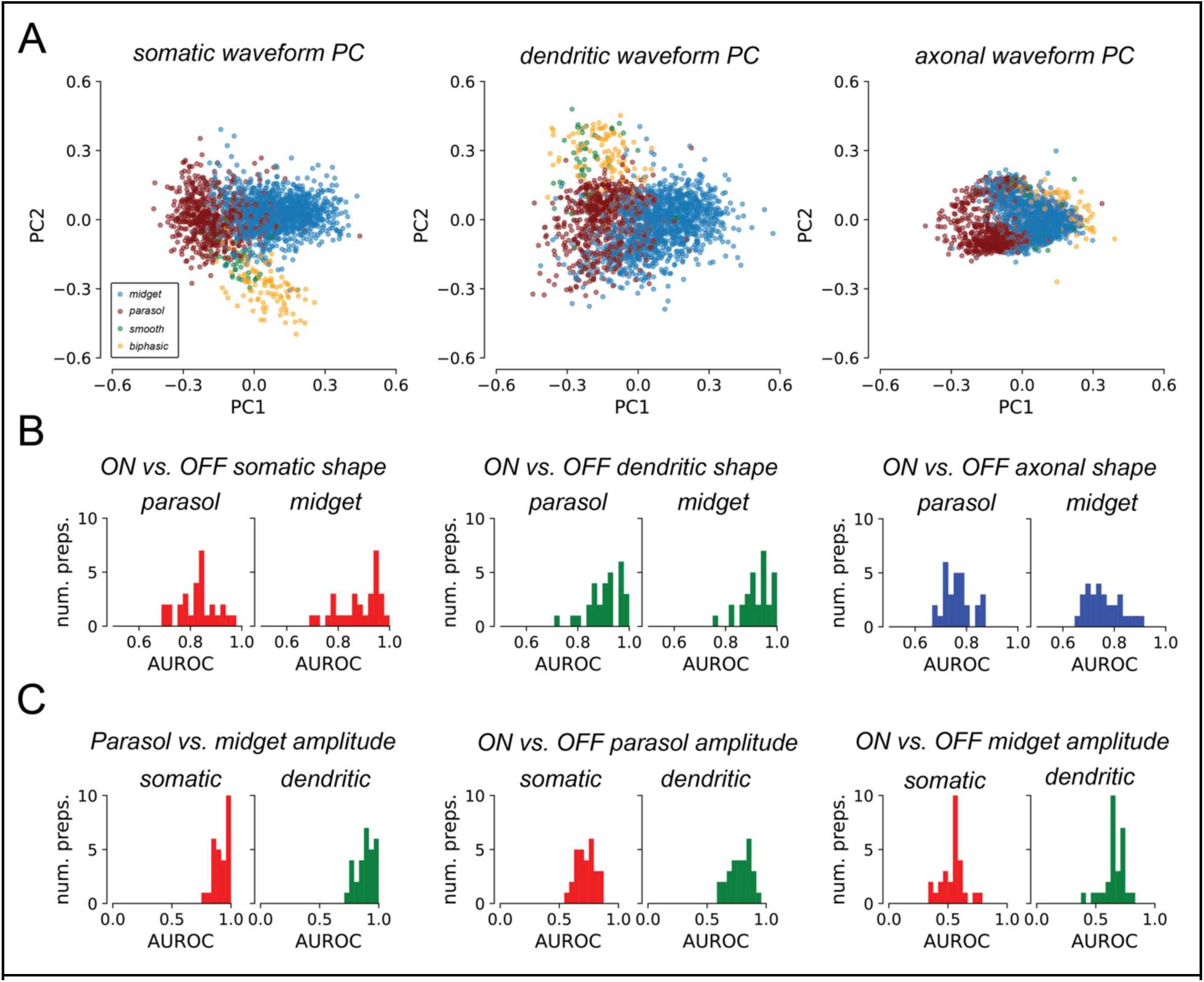
Decomposition reveals systematic differences across cell types. **(a)** Projection of the learned basis waveforms for cells from three preparations onto the first two principal components. Waveform shapes for cells of the same functional type tended to cluster together, suggesting that the learned basis waveforms capture systematic variability between different functional cell types across preparations. **(b)** AUROC histograms over 29 preparations for waveform shape-based ON vs. OFF classification, quantifying separability using logistic regression. Median AUROC was 0.836 and 0.907 for parasol and midget cells respectively with the somatic waveform, 0.909 and 0.922 with the dendritic waveform, and 0.764 and 0.747 with the axonal waveform. **(c)** AUROC histograms over 29 preparations for separability of RGC cell types by somatic and dendritic amplitude L_2_ norms. Left: Separability of parasol cells and midget cells, assuming that parasol cells have larger norms than midget cells. Median AUROC for somatic norms was 0.896, and 0.948 for dendritic norms. Middle: Separability of ON and OFF parasol cells, assuming that ON cells have larger norms than OFF cells. Median AUROC for somatic norms was 0.711, and 0.787 for dendritic norms. Right: Separability of ON and OFF midget cells assuming that ON cells have larger norms than OFF cells. Median AUROC for somatic norms was 0.549, and 0.655 for dendritic norms.

However, a similar PCA analysis failed to reveal systematic differences in waveform shape between ON and OFF RGCs across preparations (not shown). Therefore, to test whether ON and OFF RGCs could be reliably separated *within* a preparation, binary linear logistic regression classifiers were trained separately for each preparation. Classifiers for each basis waveform were fitted separately, and for parasol cells and midget cells. Separability between ON and OFF cells over the *training set* was quantified by computing the area under the receiver operator curve (AUROC), a metric for binary classifier performance (Fig. 5B). AUROC characterizes the tradeoff between classifier’s true and false positive rates, and is typically reported on a scale of 0.5 to 1, where 0.5 corresponds to random guessing and 1 corresponds to perfect classification. The median training AUROC across 29 preparations was substantially greater than 0.5 for separating ON and OFF parasol and midget cells using each of the somatic, dendritic, and axonal compartments (Fig. 5B). The clear separability reveals that the shapes of the fitted basis waveforms were informative of differences between ON and OFF RGCs.

The separability of cell types by decomposition amplitudes was explored by computing L_2_ norms for the amplitudes of each compartment of each cell, and inferring cell types using decision rules derived from anatomy. These rules were: (1) parasol RGCs have larger dendritic and somatic norms than midget RGCs, because parasol RGC dendritic trees span larger areas than those of midget RGCs [Dacey 1992, Watanabe 1989], and because parasol RGC soma diameters are larger than those of midget RGCs [Watanabe 1989]; and (2) ON cells have larger dendritic and somatic norms than OFF cells of the complementary type, because ON cells generally have have larger dendritic trees [Dacey 1992, Dacey 1993] (but see Watanabe 1989]) and larger soma diameters [Watanabe 1989] than their OFF counterparts. Though the L_2_ norm confounds the geometric size with the density and total number of ion channels, these decision rules were found to be effective in distinguishing RGCs by type.

Separability of RGC types by amplitude norms was quantified with AUROC (Fig. 5C). Though AUROC is typically reported between 0.5 and 1, AUROC for this analysis was reported between 0 and 1, as the anatomically-derived decision rules could produce worse-than-random performance. Across 29 preparations, parasol cells and midget cells were easily separated by both somatic and dendritic amplitude norms (Fig. 5C, left), in concordance with anatomy [Dacey 1992, Watanabe 1989] and prior analyses of EIs [Richard 2015]. ON and OFF parasol cells could also be separated by their somatic and dendritic norms (Fig. 5c, middle), in agreement with anatomical studies [Dacey 1992] and prior analyses of EIs [Zaidi 2023]. However, there did not appear to be systematic differences in somatic or dendritic norms between ON and OFF midget cells (Fig. 5C, right). Furthermore, the OFF midget cells in some preparations in fact had larger somatic and dendritic amplitudes than their ON counterparts. Thus the decomposition compartment amplitude norms were informative of the functional cell type of RGCs.

### Decomposition efficiently identifies RGC types in a novel retina

Finally, to demonstrate the utility of the decomposition for identifying cell types in a novel (unseen) retina, which is required for calibrating an epiretinal prosthesis, artificial neural networks were trained to use the decomposition representation alone to classify cells of the four major types across experimental preparations. The inputs to the neural networks consisted of the learned basis waveforms for each cell, and the compartment amplitude L_2_ norms z-scored within each preparation. Basis waveforms were aligned in time, but no temporal scaling was performed. Leave-one-out training and evaluation was used, wherein each instance of the neural network was trained on 28 preparations and evaluated on a single remaining heldout preparation, mimicking cell type classification in a novel retina.

Classifier performance was evaluated by computing classification accuracy and AUROC for each instance of the neural network. Metrics were computed for overall four-way classification performance, as well as for binary classification sub-problems parasol vs. midget cell, ON vs. OFF parasol cell, and ON vs. OFF midget cell. For the four-way classification problem, the classifiers achieved overall accuracy and AUROC substantially better than random chance (Fig. 6A). The decomposition featurization was sufficient for identification of parasol cells and midget cells without considering ON vs. OFF types (Fig. 6B), and for determining the ON vs. OFF cell type of corresponding RGC types (Fig. 6CD). Notably, these classifiers substantially exceeded the performance of previous EI-only approaches to this task [Zaidi 2023] (Figure 6EF) in terms of both accuracy and AUROC, and demonstrated the first high-performance EI-only method to identify ON and OFF midget RGCs (Fig. 6D). The performance of the approach did not strongly depend on the choices of amplitude L_2,1_ and waveform shape hyperparameters when fitting the decompositions (Supplemental Figure S1), showing that the decomposition effectively captures cell-type-specific properties in the EI. Thus, the decomposition representation of the EI, with no additional information, makes it possible to effectively classify and identify the major RGC types of the primate retina from electrical signals alone.

**Figure 6.**
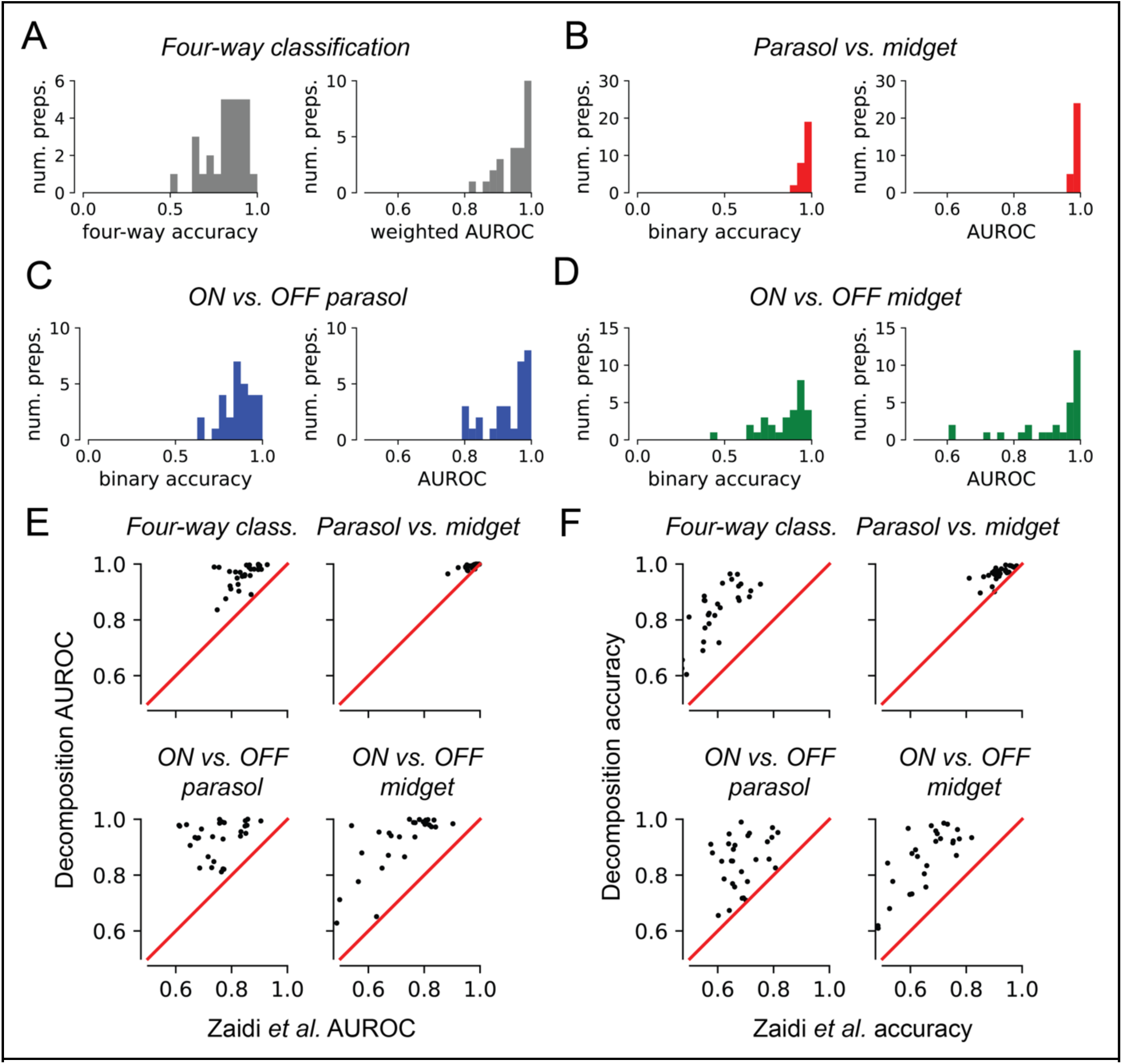
Performance of neural network cell type classifiers on held-out data. Each point represents classification performance for a single retinal recording using a neural network trained on the remaining 28 recordings. **(a)** Four-way classification performance. The median four-way accuracy was 0.850 (mean 0.825), and the median weighted one-vs-rest area under the ROC curve (AUROC) was 0.978 (mean 0.957). **(b)** Parasol vs. midget cell classification performance. Median classification accuracy was 0.977 (mean 0.966), and median AUROC was 0.994 (mean 0.991). **(c)** Classification metrics for ON vs. OFF parasol cell. Median classification accuracy was 0.867 (mean 0.855), and median AUROC was 0.959 (mean 0.932). **(d)** Classification metrics for ON vs. OFF midget cell. Median classification accuracy was 0.879 (mean 0.845), and median AUROC was 0.971 (mean 0.917). **(e)** AUROC comparison for cell-type classification, comparing decomposition-based neural network classifier performance against neural network classifiers using the EI-only features from Zaidi *et al*. [Zaidi 2023], the best-performing benchmark. For each of the classification tasks, the decomposition-based neural network achieved higher AUROC than the benchmark. **(f)** Accuracy comparison for cell-type classification, comparing decomposition-based neural network classifier performance against neural network classifiers using EI-only features [Zaidi 2023]. For each of the classification tasks, the decomposition-based neural network achieved higher accuracies than the benchmark.

The sensitivity of four-way neural network classification performance to reduced EI SNR associated with lower spike counts was evaluated in one experimental preparation. Decompositions were fitted to EIs computed with 100, 1000, and 10000 spikes, respectively. Cell type classification was performed using the appropriate instance of the previously-trained neural network. Classifier AUROC (Supplemental Fig. S3C) and accuracy (Supplemental Fig. S3D) increased systematically with increasing spike count (increasing EI SNR).

## Discussion

We have presented a method for inferring the physiological properties of neurons from recorded voltage waveforms by decomposing their electrical images (EIs) into superpositions of contributions from somatic, dendritic, and axonal compartments. We applied this technique to analyze RGCs recorded from the primate retina, demonstrating strong correlations between decomposition-fitted features and the functional, morphological, and anatomical properties of RGCs. The learned amplitudes and waveform shapes revealed systematic differences between RGCs of different functional types, enabling efficient identification of the four major RGC types in the primate retina across preparations, a substantial advance for vision restoration efforts with bi-directional implants. These findings demonstrate that electrical imaging, analyzed with suitable tools, can be a powerful tool for inference of cellular biophysics and for neuroengineering.

EI decomposition substantially advances the ability to infer the functional types of RGCs without direct characterization of their visual response properties, an important unsolved challenge in the development of bi-directional retinal implants to restore sight. Existing methods for identifying cell type without light responses [Richard 2015, Zaidi 2023] rely heavily on properties of spike timing statistics that may not be preserved during retinal degeneration because of changes in retinal circuitry or synaptic inputs [Sekirnjak 2007, Trenholm 2015, Jones 2016], and have limited ability to distinguish ON-midget from OFF-midget RGCs, the two most numerous RGC types in humans and macaques. This work demonstrates that the amplitudes and waveform shapes learned by the decomposition are each informative of the functional type of RGCs, and uses these features to train cell-type classification models that far exceed the accuracy of previously published EI-only methods [Zaidi 2023], achieving the first accurate classification of ON and OFF midget RGCs. Furthermore, as the decomposition-based classifiers used neither axon conduction velocity [Li 2012, Zaidi 2013] nor spike train statistics [Richard 2015, Zaidi 2023], features that dominated performance in past methods, it is complementary to and can be combined with those features for better overall performance.

The decomposition-based approach to RGC functional type inference may translate more effectively to the degenerated retina than spike train statistics analysis used in previous work [Richard 2015, Zaidi 2023], because RGC EIs depend primarily on intrinsic properties of the cell such as the location and density of ion channels and thus are thought to change little with degeneration (but see [Chen 2005, Chen 2013]). However, because recordings of degenerate retina were not analyzed here, quantitative evaluation of this remains a subject for future inquiry.

The performance of decomposition-based RGC functional type inference varies substantially between experimental preparations. Understanding and accounting for the underlying causes of variability between animals and preparations is an important line of future work to maximize performance in clinical applications.

The exploration of the relationship between waveform shape and functional cell type in the present work is a substantial advance over previous work using somatic spike waveform shapes to segregate cortical neurons into different categories, including coarse separation into narrow- and broad-spiking categories [Hussar 2009, Onorato 2020] and clustering for identification of potential functional types [Trainito 2019, Sun 2021, Lee 2021]. By enabling principled identification of each of the characteristic compartment waveforms and accounting for complexities in their superpositions in extracellular recordings, the decomposition extends the analysis of the relationship between waveform shape and functional cell type to the dendritic and axonal compartments, and can be used in conjunction with clustering tools to identify and characterize novel RGC types in the retina.

The EI decomposition enables more accurate inference of the spatial light response properties of RGCs without direct characterization, another fundamental computational challenge in the operation of epiretinal prosthetic devices. Decomposition-computed dendritic fields systematically reduced error in RF center estimation relative to using the inferred soma location alone, and the shapes of the decomposition dendritic fields correlated strongly with the shapes of the receptive fields. These findings could improve the accuracy of inferred RGC light response models previously proposed for determining stimulation patterns in epiretinal implants [Shah 2020, Zaidi 2023]. However, the correspondences between dendritic fields and receptive fields evaluated in this work may be incomplete because they do not take into account potential changes in receptive field shape as a function of light level.

The computational intensity of the decomposition approach significantly exceeds that of existing approaches for analyzing RGC EIs that extract simple features from recorded waveforms [Zaidi 2023], as the decomposition is computed using iterative passes of optimization and search. However, this computational intensity likely will not limit the applicability of the technique to larger datasets. This is because the decomposition is computed independently for each cell and for each recording electrode, and thus can be highly parallelized.

The decomposition fitting procedure requires that extracellular potentials from each cellular compartment of each neuron be sufficiently densely sampled to ensure reliable fitting. Recordings from two-dimensional neural tissue such as retina or neural cultures using established MEA technology [Litke 2004, Jang 2023, Zeck 2017, Müller 2015] easily meet this criterion. However, applying the decomposition method to recordings where extracellular potentials from a cell appear only on a small number of recording channels (e.g. NeuroPixel recordings from the brain [Jun 2017]) will likely prove more challenging.

Though this work demonstrates correspondence between the EI and the morphology, functional type, and visual response properties of RGCs, precise biological interpretations for the decomposition remain uncertain. The shifted superposition signal model used here assumes that each compartment basis waveform appears at most once per recording electrode, limiting the applicability in cases where cells have complicated morphologies (e.g. crossing axons in amacrine cells [Greschner 2014]) or patterns of spiking (cells that produce several distinct spike shapes, e.g. [Rhoades 2019]). Generalizations of the model will be needed to handle these cases. Furthermore, because of the difficulty of cross-validating the decomposition model and fitting hyperparameters, the amplitudes and basis waveforms learned by the decomposition cannot yet be interpreted as precise descriptions of the morphology or biophysical properties of a RGC. Additional studies combining MEA electrophysiology with higher-resolution imaging will be needed to further validate the decomposition algorithm and better understand the biophysical and morphological origins of the recorded extracellular signal. Such advances could potentially broaden the impact of electrical imaging analysis in other systems.

## Materials and Methods

### Experiments, spike sorting and EI computation

Retinas were obtained from macaque monkeys terminally anesthetized by other laboratories in the course of their experiments. All animals were handled according to approved institutional animal care and use committee (IACUC) protocols (#28860) of the Stanford University. The protocol was approved by the Administrative Panel on Laboratory Animal Care of the Stanford University (Assurance Number: A3213-01). Extracellular recording was performed using custom multielectrode arrays (MEAs) [Litke 2004], with either 512 recording electrodes with 60 um spacing and covering a 1×2mm rectangular area, or with 519 recording electrodes with 30 um spacing and covering a 1×1mm hexagonal area. Kilosort 2 was used for spike sorting [Pachitariu 2023]. RGC light response properties, including receptive fields, were characterized with reverse correlation using a white noise checkerboard stimulus [Chichilnisky 2001]. Stimuli were presented at low photopic light levels (rates of 800-2200, 800-2200, and 400-900 photoisomerizations per second for the L, M, and S cones respectively [Field 2009, Field 2010]). Ground truth cell type classification was performed manually by clustering over features computed from the light response properties and spiking auto-correlation functions, according to previously described procedures [Field 2007, Rhoades 2019]. Electrical images (EIs) for each cell were computed by cropping windows of the raw recorded voltage traces on all electrodes spanning from 3 ms (60 samples) before to 6 ms (120 samples) after each identified spike time, and then computing the mean over all windows.

### EI decomposition algorithm

The EI decomposition models each data waveform in the EI as the sum of shifted non-negative superpositions of learned compartment basis waveforms. A distinct set of basis waveforms was learned for each neuron, and that basis set is shared across every electrode of the EI for that neuron. The algorithm jointly estimated the waveforms *B*, nonnegative amplitudes *A*, and the timeshifts ***τ*** from the EI matrix *X* by approximating the solution to the constrained minimization problem

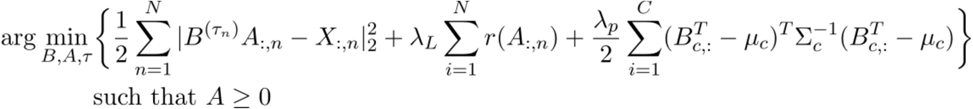

where *B^(**τ**)^* denotes shifting of the waveforms of *B* in time by ***τ**_n_* samples, *A_:,n_* denotes column *n* of the matrix *A*, *X_:,n_* denotes column *n* of matrix *X*, and *B_c,:_* denotes row *c* of matrix *B*. The first term, 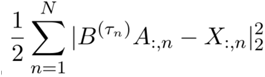, is a least squares data fidelity term. The second term, 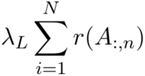, is an L_2,1_ group-sparsity-inducing regularizer on the learned amplitudes *A*. The final term, 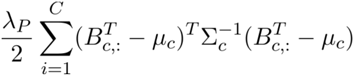, is a Gaussian prior regularizer on the waveform shapes. **λ**_L_ and **λ**_P_ are hyperparameters controlling the strength of the respective regularizers, and μ_c_ and Σ_c_^-1^ are hyperparameters describing the means and covariances of the waveform shape priors. The overall objective is non-convex, and thus fitting is performed iteratively, alternating between solving for the amplitudes and shifts assuming fixed waveform shapes, and then using those amplitudes and shifts to update the waveform shape.

The amplitudes and shifts optimization step jointly solved for the amplitudes *A* and time shifts ***τ***while holding the waveform shapes *B* fixed, resulting in the minimization problem

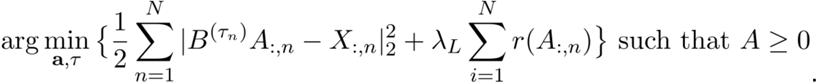

Solutions for each electrode are found independently with a coarse-to-fine search over possible combinations of time shifts ***τ**_n_*, using constrained convex minimization to find the optimal amplitudes *A_:,n_* for each possible ***τ**_n_*, and selecting the values of ***τ**_n_* and *A_:,n_* that minimize the overall objective. The L_2,1_ group-sparsity penalty used groups {soma, dendrite} and {axon}. Each constrained convex minimization problem is solved with FISTA [Beck 2009], using a modified formulation for the L_2,1_-regularized problem from [Liu 2009] that accounted for the non-negativity constraint (proof provided in the Supplement).

The waveform shape optimization step fits the waveform shapes *B* while holding the amplitudes and shifts fixed by minimizing a linear least squares objective in Fourier domain, regularized with a Gaussian prior. Letting 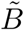 denote the discrete Fourier transform (DFT) of the basis waveforms, 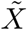 the DFT of the EI data matrix, 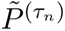 the Fourier-domain circular time shift matrix corresponding to the time shifts ***τ**_n_* found in the previous step, ⊙ Hadamard elementwise matrix multiplication, and *G* a linear transformation constructed by stacking the real and imaginary components of the DFT synthesis matrix, the waveform optimization problem is

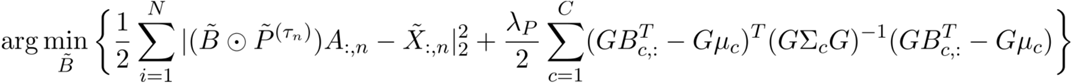

and is solved as a linear system of equations in the Fourier domain. A derivation of the coefficients of the linear system is provided in the Supplement.

The Gaussian waveform shape prior ensures that the linear system remained full-rank even if a cellular compartment is not observed. This prior term weakly constrains the basis waveform shapes to resemble the stereotyped compartment waveforms, preserving interpretability of the basis waveforms and reducing the total number of iterations required for high-quality fits.

### EI decomposition hyperparameter selection

Because of challenges in cross-validating hyperparameters, a single set of hyperparameters was selected using manual inspection of fits in one preparation. The waveform shape means μ_c_ were estimated by computing the mean waveforms for the respective compartments over all RGCs in that preparation. The waveform shape covariance matrices Σ_c_^-1^ were chosen as radial basis function kernels with length 250 μs. ⋋_*L*_, the weight placed on the sparsity regularizer, and, the weight placed on the waveform shape prior, were chosen by manual inspection of decomposition fits for that single preparation. The same hyperparameters were used for every remaining preparation, reducing the possibility of overfitting or biasing downstream analyses.

### Imaging and alignment

The alignment of decomposition somatic centers with imaged cell body locations re-analyzed the data from [Li 2014]. In brief, the location of the MEA was matched to the confocal images by manually aligning the locations of tissue landmarks over stacks of stained and labeled micrographs. An initial light-field micrograph of the retinal tissue on the MEA was taken to determine the approximate location of the tissue in relation to the recording electrodes. After recording, the retinal tissue was removed from the array, stained with βIII-tubulin to label the RGCs, and then imaged using a confocal microscope. Alignment between the brightfield image and the confocals was performed by matching tissue landmarks and computing nonlinear transformations mapping between the coordinate spaces. Because imaging required removing the retina from the MEA, this alignment could not be precisely determined due to possible warping of the retina.

### Projection targeting with phototagging in rat retina

A Long-Evans rat (Charles River) received bilateral injections into the superior colliculus with rAAVretro-CAG-ReaChR-GFP to fluorescently label RGCs on the MEA [Bohlen, 2020]. The superior colliculus was targeted using stereotaxic coordinates and verified by post-mortem histology. Six months post injection, the animal was deeply anesthetized by isoflurane and injected with ketamine (100 mg/kg; IP) and xylazine (10 mg/kg; IP), and eyes were enucleated. Next, the animal was transcardially perfused with a saline flush followed by 4% paraformaldehyde solution for brain histology. Concurrently, the eyes were hemisected, the vitreous was removed, and the posterior segment was dark adapted for at least 30 minutes at 32° C. A small sample of retina (∼2 mm x 3 mm) was then isolated and placed on the MEA. Checkerboard noise was used to elicit visual responses and measure receptive fields of RGCs under photopic conditions (∼10,000 Rh*/rod/s). A cocktail of drugs was then introduced to the bath application of Ames media including L-AP4 (100 µM, Tocris 0103), CNQX (100 µM, Tocris 1090) and DL-AP5 (100 µM, Tocris 1015), to block photoreceptor-driven responses. A 565 nm LED (Thorlabs, M565L3) was used to drive ReaChR-mediated spiking in transfected RGCs. The EI for ReaChR positive cells was used to match RGC responses pre- and post-photoreceptor block. The EI was then matched to the ReaChR-GFP-expressing RGC over the MEA, which was straightforward because RGC labeling was sparse. Following physiology, each retina was fixed using 4% PFA for 30-60 minutes and then immunolabeled for confocal imaging. The RGC was imaged with a 60x objective, reconstructed using CorelDraw, and then registered to the position of the cell on the MEA to align the cell with the EI. All procedures were approved by the Institutional Animal Care and Use Committee at Duke University and followed practices as outlined by the National Institutes of Health.

### Receptive field center estimation

Receptive field centers were computed from the spike-triggered averages (STAs) characterized with white noise reverse correlation [Chichilnisky 2001]. The time component of the STA was estimated by computing a mean over statistically-significant pixels, and a 2D intensity map was constructed by regressing the STA with that time component. Finally, the receptive field center was computed as the center-of-mass over the significant pixels in the intensity map.

### Somatic and dendritic center estimation, and coordinate transforms

Somatic centers were estimated from the decomposition by computing an amplitude-weighted center-of-mass over the recording electrode with the largest somatic amplitude, and its six nearest neighbors in the MEA hexagonal grid [Zaidi 2023]. Dendritic field centers were estimated by computing an amplitude-weighted center-of-mass over the recording electrodes that exceeded a threshold.

The somatic and dendritic components of the EI were expressed in terms of the MEA recording electrode coordinates, whereas the locations of the RGC receptive fields were expressed in terms of the coordinates of the visual stimulus. Affine mappings between electrode coordinate space and stimulus coordinate space were computed using the RANSAC algorithm [Fischler 1981], chosen for its robustness to outliers caused by the boundaries of the MEA.

### Dendritic mosaic contouring and significance testing

The spatial structure of the dendritic fields for parasol RGCs was estimated from the decomposition-computed dendritic amplitudes. Because the OFF smooth monostratified and putative broad thorny RGCs were oversplit by the spike-sorter into multiple units, each with slightly different EI [Rhoades 2019], dendritic fields were estimated by computing the maximum value of the dendritic amplitude on each recording electrode over the oversplits.

Prior to contouring, the dendritic field was first converted into a smooth 2D surface by convolving with a 2D Gaussian filter with standard deviation 54 um. The resulting dendritic field surfaces were then normalized by their respective maximum values to equalize the amplitudes between cells. Maximum-tiling dendritic field contours for each cell type were constructed by finding the threshold that maximized the uniformity index (UI), defined as the fraction of the MEA recording area contained within exactly one dendritic contour, similar to the procedure in [Gauthier 2009] for contouring RGC receptive fields.

Significance testing for dendritic spatial coordination was performed by comparing the UI of real dendritic mosaics with a null distribution constructed by randomly and independently rotating each dendritic field about its geometric centroid and recontouring (N=250). P-values were estimated as the fraction of the null distribution with a greater UI than the observed data.

### Waveform shape analysis

The waveform shape principal components analysis used three experimental preparations from different animals, each containing nearly-complete populations of the OFF parasols, OFF midgets, OFF smooth monostratified [Rhoades 2019], and putative broad thorny [Kling 2023] cells. Basis waveforms were pooled across preparations. Temporal rescaling of the waveforms was performed by equalizing the full-width half-maximum of the negative phase of the mean OFF parasol somatic waveform using b-spline interpolation and resampling. Temporal alignment was performed by aligning the principal zero-crossings for the somatic and dendritic waveforms, and by aligning the absolute minimum for the axonal waveforms. Each basis waveform was analyzed separately. The top two principal components for each basis waveform were used for visualization.

The degree to which basis waveform shapes were informative of RGC polarity was evaluated with linear logistic regression. These classifiers took a single learned compartment basis waveform as input, and predicted whether each RGC was an ON or OFF cell. Separate classifiers were trained for parasol cells and for midget cells, and every cell of the relevant types within each preparation was included in the training set for the classifiers, and the separability of cell types by waveform shape was quantified by computing AUROC over the *training* set.

### Decomposition-only cell type classifier training and evaluation

Four-layer neural networks were trained to identify RGC cell type from the decomposition. The input features consisted of the learned basis waveforms for each RGC, and the L_2_ norms of the compartment amplitudes z-scored over all included RGCs within each preparation. Basis waveforms were aligned in time without temporal rescaling. Each hidden layer contained 25 units with ReLU nonlinearities and batch normalization [Ioffe 2015]. The output layer had 4 units, with softmax activation to compute classification probabilities. Each network was trained with cross-entropy loss, using mini-batch gradient descent for 30 epochs with batch size 32 and the Adam optimizer [Kingma 2017].

Leave-one-out training and evaluation was used, using 28 of the 29 experimental preparations for training, and the one remaining preparation for evaluation. This methodology mimicked the process of classifying RGC cell type in a novel retina. Only cells belonging to the four major types were included; cells of other types and unidentified spike-sorted units were ignored.

Classification accuracy and AUROC were computed for the overall four-way classification problem, and for parasol vs. midget RGC, ON vs. OFF for parasol RGCs, and ON vs. OFF for midget RGCs. Four-way AUROC was computed as the class-weighted sum of the one-vs-rest AUROC for each cell type. For parasol vs. midget RGC classification, classification was deemed to be correct if the cell was correctly identified as a parasol RGC or a midget RGC, regardless of whether the cell was an ON or OFF cell. ON vs. OFF parasol cell classification performance was evaluated by computing the conditional probability of a cell being either an ON or OFF parasol cell given that the cell was known to be a parasol. Binary AUROC was computed by sweeping a decision threshold over the conditional probability. ON vs. OFF midget cell classification performance was characterized in a similar manner.

### Cell type classification comparison existing benchmark method

Benchmark four-layer cell type classification neural networks using the EI waveform featurization from Zaidi *et al*. [Zaidi 2023] were trained. The EI waveform features consisted of 11 amplitude and time features computed from the largest amplitude EI waveform [Zaidi 2023], and were normalized within each preparation. Each neural network hidden layer had 25 units, and the output layer had 4 units, with softmax activation to compute classification probabilities. Each network was trained with cross-entropy loss, using mini-batch gradient descent for 30 epochs with batch size 32 and the Adam optimizer [Kingma 2017]. Like for the decomposition classifier networks, leave-one-out training and evaluation was used. Performance of the decomposition networks was evaluated against the benchmark networks by comparing classification AUROC and accuracy for the four-way classification problem, for parasol vs. midget RGC, ON vs. OFF parasol RGC, and ON vs. OFF midget RGC.

### Evaluation of sensitivity to decomposition hyperparameters

Decompositions were refit for every combination of L_2,1_ and prior weight hyperparameters over a wide range of possible values on a log-spaced grid, for each of 29 experimental preparations. The sensitivity of receptive field center prediction to hyperparameters was evaluated by computing the mean prediction error across preparations for each combination of hyperparameters. The sensitivity of cell type classification performance was characterized by retraining the classification neural networks for each combination of hyperparameters, and evaluating the mean classification error and AUROC using the leave-one-out framework.

### Evaluation of sensitivity to EI SNR

The sensitivities of the cell type classification and Receptive field center prediction analyses to reduced spike count (e.g. lower signal-to-noise ratio in the EI) were evaluated in one experimental preparation. EIs for each cell were recomputed using 100, 1000, and 10000 spikes, and decompositions were refit at each spike count level with the fixed hyperparameters from previous analyses. The sensitivity of four-way major cell type classification to spike count was evaluated using the appropriate pre-trained leave-one-out neural network from the full classification analysis, and performance at each spike count level was reported using four-way AUROC and four-way classification accuracy. Receptive field center prediction using the somatic and dendritic coordinates was evaluated separately for each of the major cell types at each spike count level using the method described above. The same cells were used to perform the analyses at all spike count levels.

## Author Contributions

E.G.W. and E.J.C. conceived of the computational approach. A.M.R., M.O.B., and P.H.L. performed the experiments for labeling and tracing RGC compartments, with supervision from G.D.F. and E.J.C. A.K. and S.C. assisted with the analyses of low density primate cell types. A.M.R., P.H.L., A.K., C.R., N.B., A.R.G., N.P.S., and S.S.M. collected the electrophysiological data. A.S. and A.M.L. provided the recording electronics. E.G.W. and E.J.C. wrote the manuscript, with contributions from A.M.R., M.B., and G.D.F.

## Acknowledgements

E.G.W. was supported by a National Defense Science and Engineering Graduate (NDSEG) Fellowship. E.J.C. was supported by Wu Tsai Neurosciences Institute Big Ideas, NEI grants R01EY017992 and R01EY029247. G.D.F. was supported by NEI grant R01EY034004.

