## Supplementary Material for "Decomposition of retinal ganglion cell electrical images for cell type and functional inference"

### Supplementary Information

### Supplementary Figures

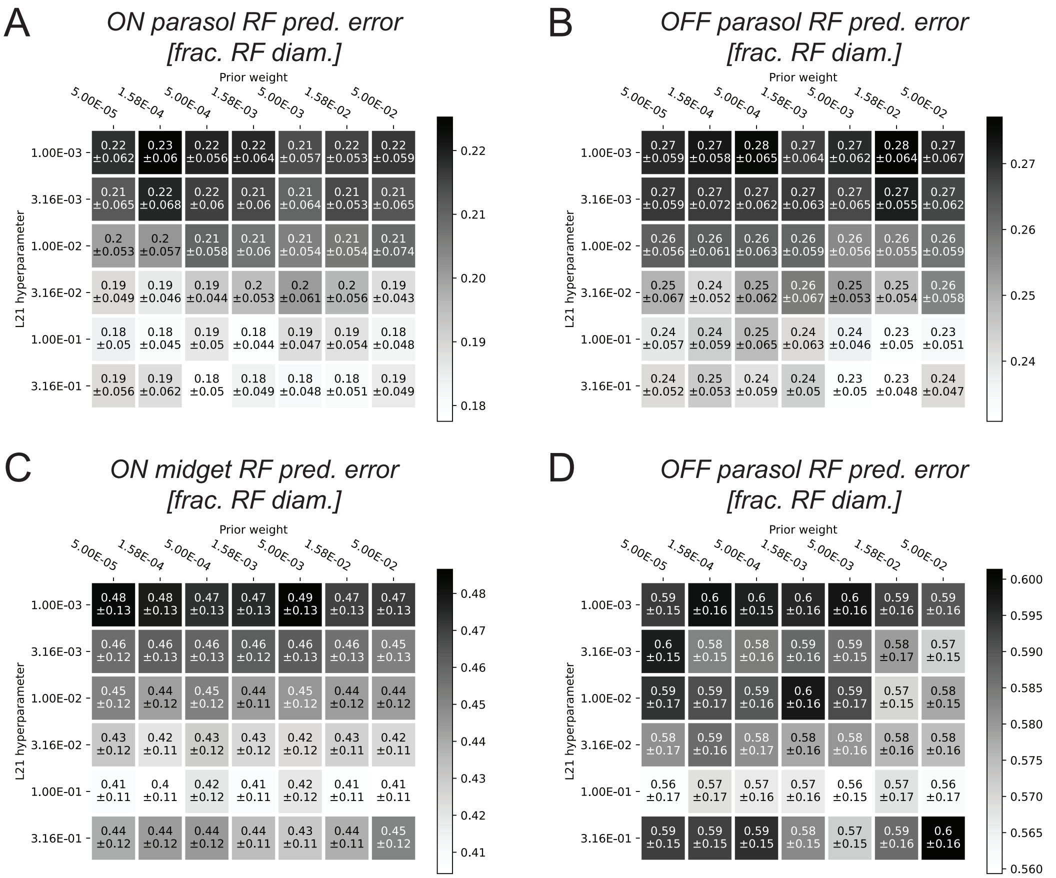


**Figure S1:** Sensitivity of major cell type receptive field center estimation to choice of decomposition hyperparameters. The EI decomposition for every cell in 29 retinas was recomputed for every combination of L2,1 and prior weight hyperparameters over a grid. Receptive field centers were estimated as an affine transform of the decomposition-estimated somatic and dendritic centers. As in Figure 3, affine transforms were fitted separately for each cell type in each retina to align electrical data to the visual stimulus. **(a)** Mean ON parasol cell receptive field center prediction error and standard deviation, expressed in units of receptive field diameter, for every combination of decomposition hyperparameters. The mean is computed as the average estimation error over all retinas. **(b)** Same as (a), for OFF parasol cell receptive field center prediction. **(c)** Same as (a), for ON midget cell receptive field center prediction. **(d)** Same as (a), for OFF midget cell receptive field center prediction. In all cases, the receptive field estimation error only weakly depends on the choice of decomposition hyperparameters.

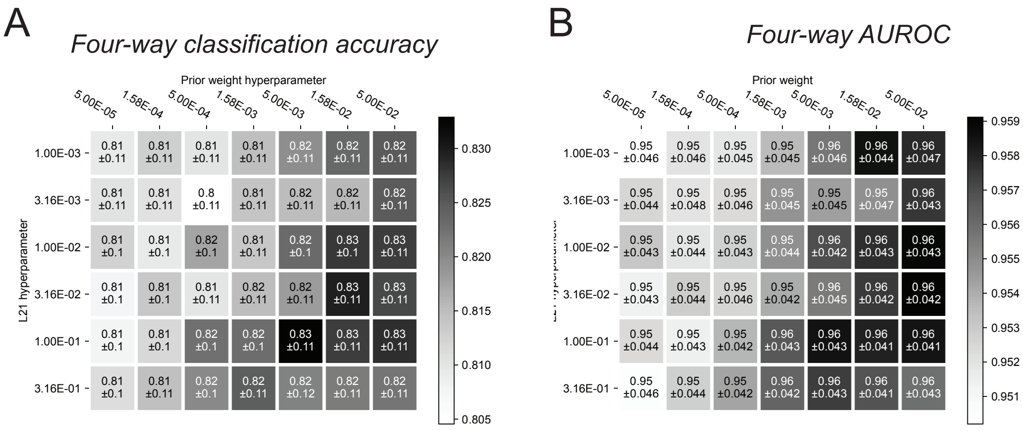


**Figure S2:** Sensitivity of major cell type classification to choice of decomposition hyperparameters. The neural network cell type classification analysis from Figure 6 was repeated over a grid of possible decomposition L2,1 and prior weight values. Evaluation for each combination of hyperparameters was performed using leave-one-out evaluation with 29 retinas. **(a)** Mean four-way cell type classification accuracy and standard deviation for every combination of decomposition hyperparameters. **(b)** Mean four-way AUROC and standard deviation for every combination of decomposition hyperparameters. In both cases, the classification AUROC did not depend strongly on the choice of decomposition hyperparameters, demonstrating the decomposition robustly extracts cell-type-specific features over a large range of possible hyperparameters.

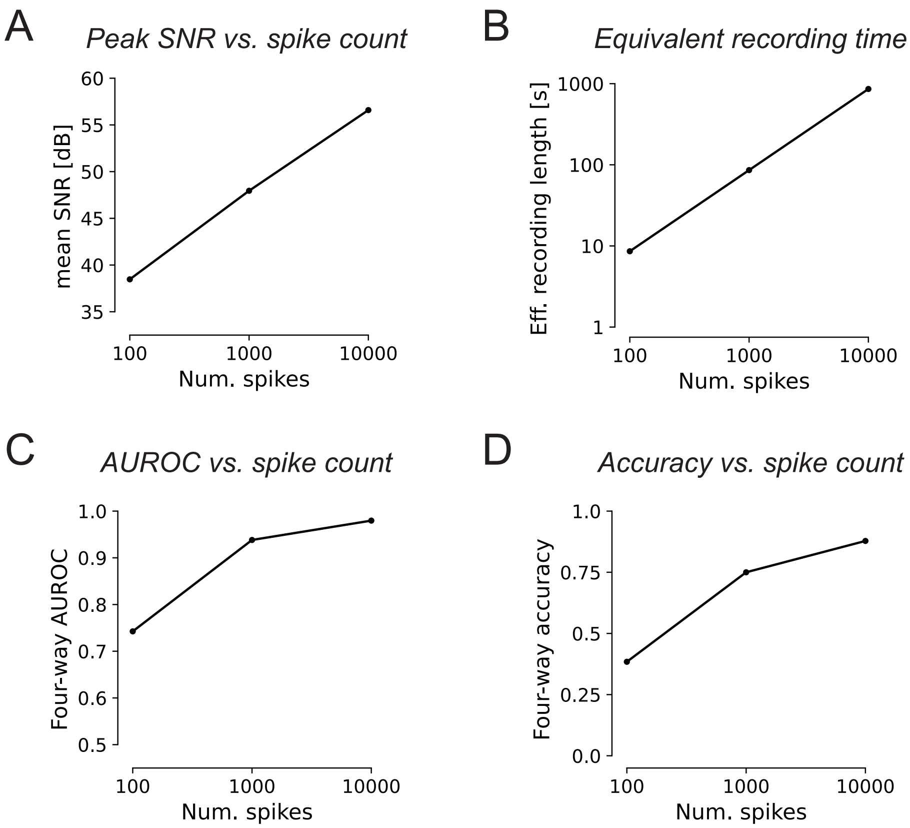


**Figure S3:** Sensitivity of decomposition-based major cell type classification performance to noise in the EI, evaluated using one experimental preparation. EIs for each RGC were computed using 100, 1000, and 10000 spikes, and performance of the pre-trained neural network classifier was evaluated at each spike count level. **(a)** Peak signal-to-noise ratio (SNR) in the EI averaged across cells at each spike count level. The peak magnitude signal was defined as the maximum deviation from zero of the recording channel with the largest amplitude signal. **(b)** Effective recording length for each spike count level, defined as the mean recording time required to observe the desired number of spikes. The 100 and 1000 spike count levels corresponded to very short equivalent recordings of length 10 and 100 seconds, respectively. **(c)** Four-way classifier AUROC for major cell type classification at each spike count level. AUROC increased with increasing spike count (increasing SNR), suggesting a strong dependence between classification performance and spike count. **(d)** Four-way classification accuracy for major cell type classification at each spike count level. Accuracy increased with increasing spike count (increasing SNR), suggesting a strong dependence between classification performance and spike count.

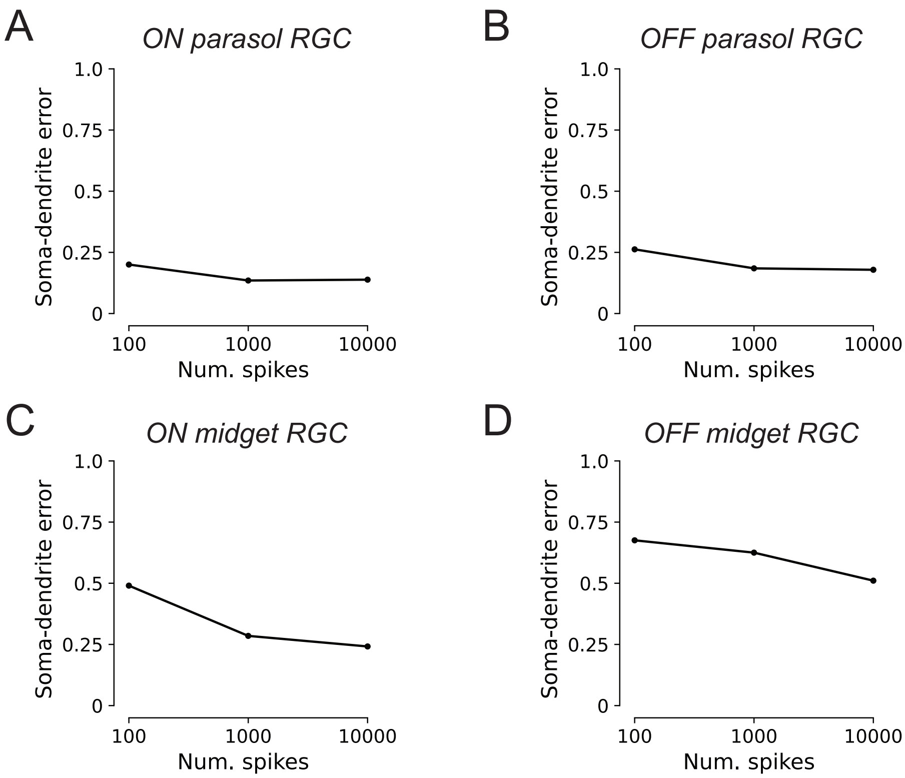


**Figure S4:** Sensitivity of receptive center estimation to noise in the EI using somatic and dendritic coordinates, evaluated in one experimental preparation. Error in each panel (y-axis) is reported in terms of fractions of receptive field diameters. **(a)** Performance of receptive field center estimation for ON parasol RGCs as a function of the number of spikes used to compute the EI. **(b)** Same as (a), for OFF parasol RGCs. **(c)** Same as (a), for ON midget RGCs. **(d)** Same as (a), for OFF midget RGCs. Performance in OFF midget RGCs was largely unaffected. Except for ON midget cells with 100 spikes, performance was also largely unaffected by the number of spikes used to compute the EI, indicating robustness to reduced EI SNR.

#### EI Decomposition Algorithm

#### 2.1 Fitting the EI decomposition

##### 2.1.1 Definitions

- The matrix $X \in \mathbb{R}^{T \times N}$ represents the EI of a neuron. This matrix contains $N$ observed voltage waveforms, one for each of the $N$ recording electrodes. Each observed waveform has temporal length $T$.
- The decomposition is parameterized by the following variables:
  - $B \in \mathbb{R}^{C \times T}$, the basis waveforms matrix, consisting of $C$ distinct basis waveforms, each with temporal length $T$.
  - $A \in \mathbb{R}^{C \times N}_{\geq 0}$, the learned amplitudes (weights) on each of the basis waveforms for every recording electrode. Every element in $A$ is required to be non-negative, because the amplitudes are interpreted as the contributions of physical compartments of the neuron to the overall recorded signal.
  - $\tau \in \mathbb{Z}^{C \times N}$, the integer-valued time shifts applied to each basis waveforms for every recording electrode.

##### 2.1.2 Decomposition fitting algorithm

The EI decomposition is fitted by minimizing the mean square error between the recorded waveforms of the EI and the waveforms reconstructed from the decomposition parameters, with additional regularization terms to induce sparsity in the learned weights and to weakly constrain the compartment basis waveform shape. Let $B^{(\tau_n)}$ denote temporal shifting of each respective waveform in $B$ by the numbers of samples specified by column $n$ of the matrix $\tau$, $A_{:,n}$ denote column $n$ of the matrix $A$, $X_{:,n}$ denote column $n$ of the matrix $X$, and $B_{:,c}$ denote row $c$ of the matrix $B$. Fitting the EI decomposition corresponds to the solving the optimization problem

$$\arg\min_{B,A,\tau} \left\{ \frac{1}{2} \sum_{n=1}^{N} |B^{(\tau_n)} A_{:,n} - X_{:,n}|_2^2 + \lambda_L \sum_{i=1}^{N} r(A_{:,n}) + \frac{\lambda_p}{2} \sum_{i=1}^{C} (B_{c,:}^T - \mu_c)^T \Sigma_c^{-1} (B_{c,:}^T - \mu_c) \right\}. \quad (2.1)$$
$$\text{such that } A \geq 0$$

The first term in the objective, $\frac{1}{2}\sum_{n=1}^{N} |B^{(\tau_n)} A_{:,n} - X_{:,n}|_2^2$, is a least squares error term, ensuring that the decomposition accurately represents the EI. The second term, $\lambda_L \sum_{i=1}^{N} r(A_{:,n})$, is a sparsity regularizer applied to the learned weights, and has associated strength hyperparameter $\lambda_L$. $r(A_{:,n})$ is specifically chosen as an $L_{2,1}$ group-sparsity regularizer, with groups {soma, dendrite} and {axon} to allow the somatic and dendritic components to appear simultaneously on the same electrode. The final term, $\frac{\lambda_p}{2}\sum_{c=1}^{C}(B_{c,:}^T - \mu_c)^T \Sigma_c^{-1}(B_{c,:}^T - \mu_c)$, is a Gaussian prior regularizer used to control the shapes of each of the learned basis waveforms, and has associated strength hyperparameter $\lambda_p$, as well as hyperparameters $\mu_c$ and $\Sigma_c$ specifying the means and covariances for each basis waveform.

The overall optimization problem (2.1) is not jointly convex in terms of the decomposition model parameters $\{B, A, \tau\}$. This is both because $\tau$ is integer-valued, and because of the simultaneous estimation of $B$ and $A$. Due to this non-convexity, the decomposition is fitted using a novel iterative algorithm for shifted semi-non-negative matrix factorization. The algorithm alternates between two steps: (1) an amplitude and time shift optimization step, holding the compartment basis waveforms fixed; and (2) a basis waveform shape fitting step, using the previously-inferred amplitudes and time shifts to learn the waveform shape. Pseudocode for the overall fitting algorithm is provided in Algorithm 1, and mathematical details for each of the sub-steps are provided subsequently.

**Algorithm 1** Alternating optimization for EI decomposition fitting

Hyperparameters: $\lambda_L$, $\lambda_p$, $\Sigma_i$, $\mu_i$
**procedure** EI-DECOMPOSITION($X$)
  Initialize $\mathbf{B}^{(k)}$ to the mean waveform values
  **for** $k \in 1, 2, \ldots, K$ **do**
    $A^{(k)}, \tau^{(k)} \leftarrow \arg\min_{A,\tau} \left\{ \frac{1}{2} \sum_{n=1}^{N} |[B^{(k)}]^{(\tau_n^{(k)})} A_{:,n} - \mathbf{x}_n|_2^2 + \lambda_L \sum_{n=1}^{N} r(A_{:,n}) \right\},\ A \geq 0$
    $B^{(k)} \leftarrow \arg\min_{B} \left\{ \frac{1}{2} \sum_{n=1}^{N} |B^{(\tau_n^{(k)})} A_{:,n} - \mathbf{x}_n|_2^2 + \frac{\lambda_p}{2} \sum_{c=1}^{C} (B_{c,:}^T - \mu_c)^T \Sigma_c^{-1} (B_{c,:}^T - \mu_c) \right\}$
    Normalize rows of $B^{(k)}$ to have $L_2$-norm 1.
  **end for**
  **return** $B, \mathbf{a}, \tau$
**end procedure**

**Amplitudes and time shifts optimization**

The amplitude and time shift optimization step holds the compartment basis waveforms $B$ fixed, and uses a combination of search and convex minimization to simultaneously fit the amplitudes $A$ and time shifts $\tau$ on every electrode. The overall objective for this step is constructed by discarding all terms that depended solely on $B$ from the objective of problem (2.1), leaving the optimization problem

$$\arg\min_{A,\tau} \left\{ \frac{1}{2} \sum_{n=1}^{N} |B^{(\tau_n)} A_{:,n} - X_{:,n}|_2^2 + \lambda_L \sum_{i=1}^{N} r(A_{:,n}) \right\} \text{ such that } A \geq 0. \tag{2.2}$$

Though problem (2.2) is still not jointly convex in terms of $A$ and $\tau$, its structure is considerably simpler than that of the full problem (2.1). In particular, problem (2.2) is separable by electrode and can be solved separately for each electrode. In addition, if the time shifts $\tau$ are held fixed, each single-electrode problem simplifies into a regularized non-negative linear least square problem,

$$\arg\min_{A_{:,n}} \left\{ \frac{1}{2} |B^{(\tau_n)} A_{:,n} - X_{:,n}|_2^2 + \lambda_L r(A_{:,n}) \right\} \text{ such that } A_{:,n} \succeq \mathbf{0}, \tag{2.3}$$

which is a constrained convex minimization problem. Leveraging this property, the amplitudes and time shifts optimization is performed using a coarse-to-fine search over candidate time shifts, solving problem (2.3) with convex minimization for each candidate time shift, and then selecting the combination of $\{A_{:,n}, \tau_n\}$ that minimizes the objective. Because the optimization is solved separately for each recording electrode, this approach avoids combinatorial explosion over the possible combinations of time shifts. Each convex minimization problem is solved using FISTA [1], an accelerated proximal gradient descent algorithm, using a modified form of the constrained optimization formulation of the $L_{2,1}$-regularized problem described in [2] to support the additional non-negativity constraint on the learned weights. A detailed description as well as mathematical proof for this formulation is provided in Section 3.1.2.

Pseudocode for the amplitudes and time shifts optimization is presented in Algorithm 2. Notably, the optimization problems within both the coarse and fine searches can be easily parallelized for improved runtime.

**Algorithm 2** Amplitudes and time shifts coarse-to-fine search

Hyperparameters: $\lambda_L$
**procedure** AMPLITUDE-TIME-SEARCH($X$, $B$)
  **for** electrode $n \in 1, 2, \ldots, N$ **do**
    $\hat{V}_n \leftarrow \infty$
    $\hat{A}_{:,n}, \hat{\tau}_n \leftarrow$ empty values
    **for** $\tau'_n \in$ coarse grid of possible time shifts **do** ▷ Coarse search over $\tau$
      $A'_{:,n} \leftarrow \arg\min_{\mathbf{a}_n} \{\frac{1}{2}|B^{(\tau_n)}A_{:,n} - X_{:,n}|_2^2 + \lambda_L r(A_{:,n})\}$ such that $A_{:,n} \succeq \mathbf{0}$
      $V'_n \leftarrow \frac{1}{2}|B^{(\tau'_n)}A'_{:,n} - X_{:,n}|_2^2 + \lambda_L r(A'_{:,n})$
      **if** $V'_n < \hat{V}_n$ **then**
        $\hat{V}_n \leftarrow V'_n$
        $\hat{A}_{:,n}, \hat{\tau}_n \leftarrow A'_{:,n}, \tau'_n$
      **end if**
    **end for**
    $V^*_n \leftarrow \infty$
    $A^*_{:,n}, \tau^*_n \leftarrow$ empty values
    **for** $\tau'_n in$ fine grid of time shifts in neighborhood of $\hat{\tau}_n$ **do** ▷ Fine search over $\tau$
      $A'_{:,n} \leftarrow \arg\min_{A_{:,n}} \{\frac{1}{2}|B^{(\tau_n)}A_{:,n} - X_{:,n}|_2^2 + \lambda_L r(A_{:,n})\}$ such that $A_{:,n} \succeq \mathbf{0}$
      $V'_n \leftarrow \frac{1}{2}|B^{(\tau'_n)}A'_{:,n} - X_{:,n}|_2^2 + \lambda_L r(A'_{:,n})$
      **if** $V'_n < V^*_n$ **then**
        $V^*_n \leftarrow V'_n$
        $A^*_{:,n}, \tau^*_n \leftarrow \mathbf{a}'_n, \tau'_n$
      **end if**
    **end for**
  **end for**
  **return** $A^*, T^*$
**end procedure**

##### Compartment basis waveform shape optimization

The compartment basis waveform shape optimization step holds the amplitudes $A$ and time shifts $\tau$ fixed, and solves the convex quadratic minimization problem

$$\arg\min_{B} \Big\{ \frac{1}{2} \sum_{n=1}^{N} |B^{(\tau_n)}\mathbf{a}_n - \mathbf{x}_n|_2^2 + \frac{\lambda_P}{2} \sum_{c=1}^{C} (\mathbf{b}_c - \mu_c)^T \Sigma_C^{-1} (\mathbf{b}_c - \mu_c) \Big\} \tag{2.4}$$

for the basis waveform shapes $B$. Construction of the matrix $B^{(\tau_n)}$ requires temporal shifting of the basis waveforms comprising $B$, complicating computer representations of problem (2.4). Because a circular shift of $t$ samples in time domain can be expressed simply in Fourier domain as multiplication by the complex phase shift $e^{-i\omega t}$, the difficulty of representing time shifts is alleviated by solving the problem in Fourier domain. Letting $\tilde{B}$ denote the discrete Fourier transform of the basis waveforms, $\tilde{X}$ the discrete Fourier transform of the observed EI, $\odot$ Hadamard (element-wise) matrix multiplication, $\tilde{P}^{(\tau_n)}$ the complex-valued Fourier phase shift matrix corresponding to temporally shifting the compartment basis waveforms by the amounts specified in $\tau_n$, and $G$ the stacked real-imaginary discrete Fourier transform analysis matrix (see Section 4.1.2 for definition), problem (2.4) is expressed in Fourier domain as

$$\arg\min_{\tilde{B}} \left\{ \frac{1}{2} \sum_{i=1}^{N} |(\tilde{B} \odot \tilde{P}^{(\tau_n)})A_{:,n} - \tilde{X}_{:,n}|_2^2 + \frac{\lambda_P}{2} \sum_{c=1}^{C} (GB_{c,:}^T - G\mu_c)^T (G\Sigma_c G)^{-1} (GB_{c,:}^T - G\mu_c) \right\}. \tag{2.5}$$

This formulation is equivalent to the original time domain problem (2.4) due to Parseval's relation. As problem (2.5) corresponds to minimization of a convex quadratic, it is solved by constructing and solving a system of linear equations derived from the first-order conditions of optimality. A complete derivation of the coefficients of this linear system is provided in Section 4.1.

#### Proximal Gradient Descent Algorithms

#### 3.1 Non-negative $L_{2,1}$-regularized minimization

The EI decomposition fitting procedure solves problem (2.3) as a non-negative $L_{2,1}$-regularized linear least squares problem. Because the $L_{2,1}$ regularizer is not smooth, this problem cannot be solved directly with gradient descent. Following [2], we reformulate the problem into a different constrained convex minimization problem that can be easily solved using proximal gradient descent. The derivation of the proximal operator for the re-formulated problem builds heavily on the original derivation of the unconstrained $L_{2,1}$-regularized proximal operator performed in [2], and so we repeat that derivation first.

##### 3.1.1 Unconstrained $L_{2,1}$-regularized minimization

*The derivation for the proximal algorithm solving the unconstrained $L_{2,1}$-regularized convex minimization problem (e.g. without the non-negativity constraint) is taken from [2].*

Consider the *unconstrained* $L_{2,1}$-regularized convex minimization problem

$$\min_{W} \left\{ L(W) + \rho |W|_{2,1} \right\} \tag{3.1}$$

where $L(W)$ is a smooth convex loss function in the variable $W$. This problem cannot be solved using vanilla gradient descent, because the $L_{2,1}$ penalty term $|W|_{2,1}$ is not differentiable everywhere (e.g. at the origin). Reference [2] replaces this unconstrained non-smooth problem with the constrained smooth optimization problem

$$\min_{(W,\mathbf{t}) \in D} \left\{ L(W) + \rho \sum_{i=1}^{G} t_i \right\} \tag{3.2}$$

where $D$ is the convex set $D = \{(W, \mathbf{t}) \mid |\mathbf{w}_i|_2 \leq t_i,\ i \in 1, 2, ..., G\}$ and $G$ is the number of groups in the $L_{2,1}$ group-sparsity regularizer. By inspection, $D$ is convex, since it is the Cartesian product of ice-cream-cones. The objective in problem (3.2) is smooth and convex, and thus problem (3.2) is a constrained smooth convex minimization problem. This problem can be solved using proximal gradient descent or one of its accelerated variants (e.g. FISTA [1]), provided that projection onto the convex set $D$ can be performed efficiently. This projection problem, projecting an arbitrary $(U, \mathbf{v})$ onto the convex set $D$, is expressed as

$$\pi_D(W, \mathbf{t}) = \arg \min_{(U,\mathbf{v}) \in D} \left\{ \frac{1}{2} |W - U|_F^2 + \frac{1}{2} |\mathbf{t} - \mathbf{v}|_2^2 \right\}. \tag{3.3}$$

Equation (3.3) can be separated by group, resulting in

$$\pi_D(W, \mathbf{t}) = \arg \min_{(U,\mathbf{v}) \in D} \left\{ \sum_{i=1}^{G} \left[ \frac{1}{2} |\mathbf{w}_i - \mathbf{u}_i|^2 + \frac{1}{2} (t_i - v_i)^2 \right] \right\}. \tag{3.4}$$

Assuming that none of the variables in $W$ are shared between groups, problem (3.4) is solved found by separately solving each of the one-group problems

$$\pi_{D_i}(\mathbf{w}_i, t_i) = \arg \min_{(\mathbf{u}_i, v_i) \in D} \left\{ \frac{1}{2} |\mathbf{w}_i - \mathbf{u}_i|^2 + \frac{1}{2} (t_i - v_i)^2 \right\}. \tag{3.5}$$

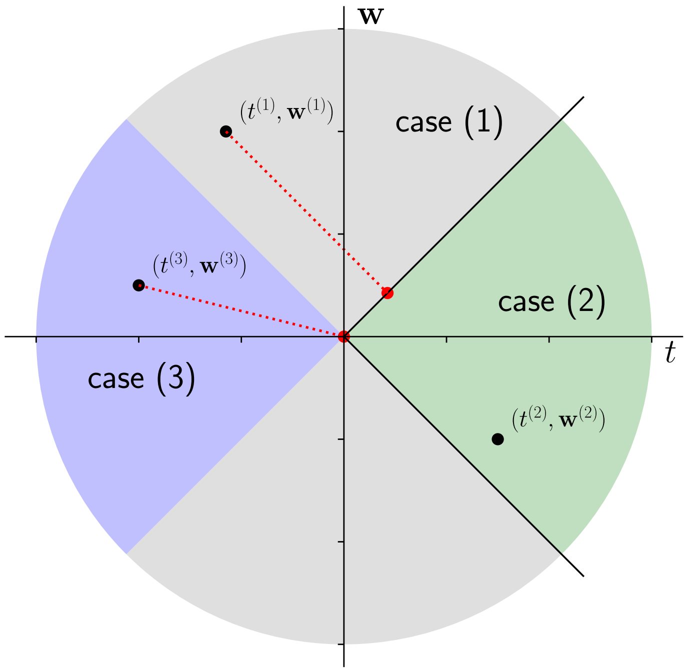


**Figure S1:** Projection onto the 1D ice-cream cone. There are three possible cases: (1) $|w_i| > |t_i|$, corresponding to the gray shaded region; (2) $|w_i| \leq |t_i|$, corresponding to the green shaded feasible region; (3) $|w_i| \leq -t_i$, corresponding to the blue shaded region. In cases (1) and (3), $(w_i, t_i)$ lies outside the feasible region, and the associated dotted red lines and points correspond to the minimum norm projections onto the feasible set.

Consider the case where $\mathbf{w}_i$ is one-dimensional. A drawing of the corresponding two-dimensional $(\mathbf{w}_i, t_i)$ coordinate space illustrating the three possible cases for projection is provided in Figure S1. The first case, $|w| > |t|$, corresponds to the gray shaded region of Figure S1. This is outside the feasible region, and the projection corresponds the orthogonal projection onto the edge of the ice-cream cone. The second case, $|w| \leq |t|$, corresponds to the green shaded region of Figure S1. This is already feasible, so nothing needs to be done. The third case, $|w| \leq -t$, corresponds to the blue shaded region of Figure S1. This is outside the feasible region, and the solution is to project onto the origin. Formally, these rules can be expressed as

$$u_i^* = \begin{cases} \frac{w_i + t_i}{2}, \ |w_i| > |t_i| & \text{(case 1)} \\ w_i, \ |w_i| \leq t_i & \text{(case 2)} \\ 0, \ |w_i| \leq -t_i & \text{(case 3)} \end{cases}$$

$$v_i^* = \begin{cases} \frac{w_i + t_i}{2}, \ |w_i| > |t_i| & \text{(case 1)} \\ v_i, \ |w_i| \leq t_i & \text{(case 2)} \\ 0, \ |w_i| \leq -t_i & \text{(case 3)} \end{cases}$$

Because the cone is rotationally-symmetric about the $t$ axis, the solution to the one-dimensional projection problem can be generalized to $N$ dimensions as

$$\mathbf{u}_i^* = \begin{cases} \frac{|\mathbf{w}_i|+t_i}{2}\frac{\mathbf{w}_i}{|\mathbf{w}_i|},\ |\mathbf{w}_i| > |t_i| & \text{(case 1)} \\ \mathbf{w}_i,\ |\mathbf{w}_i| \le t_i & \text{(case 2)} \\ \mathbf{0},\ |\mathbf{w}_i| \le -t_i & \text{(case 3)} \end{cases} \tag{3.6}$$

$$t_i^* = \begin{cases} \frac{|\mathbf{w}_i|+t_i}{2},\ |\mathbf{w}_i| > |t_i| & \text{(case 1)} \\ t_i,\ |\mathbf{w}_i| \le t_i & \text{(case 2)} \\ 0,\ |\mathbf{w}_i| \le -t_i & \text{(case 3)} \end{cases} \tag{3.7}$$

*Proof.* (3.6) and (3.7) solve the optimization problem (3.5).

The proof is performed by verifying that (3.6) and (3.7) satisfy the following condition of optimality for constrained smooth convex minimization problems: *For the convex minimization problem* $\arg\min_{\mathbf{x}\in F}\{L(\mathbf{x})\}$ *with smooth convex objective* $L(\mathbf{x})$ *and convex feasible set* $F$*, the point* $\mathbf{x}^* \in F$ *is optimal if and only if* $\langle \mathbf{x}-\mathbf{x}^*, L'(\mathbf{x}^*)\rangle \ge 0$*,* $\forall \mathbf{x} \in F$. The gradient of the objective in (3.5) with respect to $(\mathbf{u}_i, v_i)$ is $(\mathbf{u}_i - \mathbf{w}_i, v_i - t_i)$. Consider each of the three possible cases:

- Case (1), $|\mathbf{w}_i| > t_i$. Define $\alpha = \frac{t_i - |\mathbf{w}_i|}{2}\frac{\mathbf{w}_i}{|\mathbf{w}_i|}$ and $\beta = \frac{|w_i| - t_i}{2}$. By inspection, $|\alpha| = \beta > 0$, where the inequality is satisfied due to the condition.

  Plugging this into the left hand side of the optimality condition, we get

$$\begin{aligned} \langle \mathbf{u}_i - \mathbf{u}_i^*, \mathbf{u_i}^* - \mathbf{w}_i\rangle + (v_i - v_i^*)(v_i^* - t_i) &= \langle \mathbf{u}_i, \alpha\rangle + t_i\beta \\ &\ge -|\mathbf{u}_i||\alpha| + t_i\beta \\ &\ge -t_i\beta + t_i\beta \\ &\ge 0 \end{aligned}$$

  where the first inequality is due to Cauchy-Schwarz, and the second inequality is due to $|\mathbf{u}_i| < v_i$, which results from the constraint $(\mathbf{u}_i, v_i) \in D$. Thus case (1) of (3.6) and (3.7) is proven.

- Case (2), $|\mathbf{w}_i| \le t_i$. In this case, $(\mathbf{u}_i^*, v_i^*) = (\mathbf{w}_i, t_i)$ is in the feasible set and minimizes the single-group objective function (3.5). Thus case (2) of (3.6) and (3.7) is proven.

- Case (3), $|\mathbf{w}_i| \le -t_i$. we write

$$\begin{aligned} \langle \mathbf{u}_i - \mathbf{u}_i^*, \mathbf{u_i}^* - \mathbf{w}_i\rangle + (v_i - v_i^*)(v_i^* - t_i) &= \langle \mathbf{u}_i, -\mathbf{w}_i\rangle + (v_i)(-t_i) \\ &\ge -|\mathbf{u}_i||\mathbf{w}_i| - t_i v_i \\ &\ge -t_i(-v_i) - t_i v_i \\ &\ge 0 \end{aligned}$$

  where the first inequality is due to the Cauchy-Schwarz inequality, and the second inequality comes from the conditions $|\mathbf{w}_i| \le -t_i$ and $t_i \le 0$. Thus case (3) of equations (3.6) and (3.7) is proven.

Thus the claim is proven. □

The overall FISTA algorithm solving the unconstrained $L_{2,1}$-regularized problem as a constrained convex minimization problem is given in Algorithm 3. Note that $f_{\gamma,W_i,\mathbf{t}_i}(W_{i+1}, \mathbf{t}_{i+1})$ in the backtracking line search is defined as

$$\begin{aligned} f_{\gamma,W_i,\mathbf{t}_i}(W,\mathbf{t}) \triangleq f(W_i,\mathbf{t}_i) + \langle \boldsymbol{\nabla}'_W f(W',\mathbf{t}')|_{W'=W_i,\mathbf{t}'=\mathbf{t}_i}, W - W_i\rangle + \\ \langle \boldsymbol{\nabla}_{\mathbf{t}'} f(W',\mathbf{t}')|_{W'=W_i,\mathbf{t}'=\mathbf{t}_i}, \mathbf{t} - \mathbf{t}_i\rangle + \frac{\gamma}{2}|W - W_i|_F^2 + \frac{\gamma}{2}|\mathbf{t} - \mathbf{t}_i|_2^2 \end{aligned} \tag{3.8}$$

and $\pi_{D_G}$ corresponds to the projection operator from equation (3.3).

**Algorithm 3** FISTA for unconstrained $L_{2,1}$-regularized problem

Hyperparameter $\rho_{L21}$
**procedure** UNCONSTRAINED-FISTA-L21($L(W)$, $W_0$, $G$, $\gamma$)
  Randomly initialize variable $\mathbf{t} \in \mathbb{R}^{|G|}_{\geq 0}$
  $f(W, \mathbf{t}) \triangleq L(W) + \rho_{L21} \mathbf{1}^T \mathbf{t}$
  Initialize $i = 1, W_1 = W_0, b_{-1} = 0, b_0 = 1$
  $W^\star = W_0$
  **repeat**
    $\alpha_i = \frac{b_{i-2}-1}{b_{i-1}}$
    $W_i' = W_i + \alpha_i(W_i - W_{i-1})$
    $\mathbf{t}_i' = \mathbf{t}_i + \alpha_i(\mathbf{t}_i - \mathbf{t}_{i-1})$
    **while** True **do** ▷ Backtracking line search
      $\gamma = \frac{\gamma}{2}$
      $\bar{W}_{i+1} = W_i' - \frac{1}{\gamma} \boldsymbol{\nabla}_W f(W, \mathbf{t})\big|_{W=W_i', \mathbf{t}=\mathbf{t}_i'}$
      $\bar{\mathbf{t}}_{i+1} = \mathbf{t}_i' - \frac{1}{\gamma} \boldsymbol{\nabla}_\mathbf{t} f(W, \mathbf{t})\big|_{W=W_i', \mathbf{t}=\mathbf{t}_i'}$
      $W_{i+1}, \mathbf{t}_{i+1} = \pi_{D_G}(\bar{W}_{i+1}, \bar{\mathbf{t}}_{i+1})$
      **if** $f(W_{i+1}, \mathbf{t}_{i+1}) \leq f_{\gamma, W_i, \mathbf{t}_i}(W_{i+1}, \mathbf{t}_{i+1})$ **then**
        $W^\star = W_{i+1}$
        break
      **end if**
    **end while**
    $b_i = \frac{1+\sqrt{1+4b_{i-1}^2}}{2}$
  **until** Converged
  **return** $W^\star$
**end procedure**

##### 3.1.2 $L_{2,1}$-regularized minimization with non-negativity constraint

*This derivation is performed by the authors.*

For the EI decomposition, the learned compartment amplitudes are required to be non-negative, corresponding to the constrained minimization problem

$$\arg\min_{W \geq 0} \left\{ L(W) + \rho |W|_{2,1} \right\} \tag{3.9}$$

where $W \geq 0$ constrains all variables in $W$ to be non-negative, $L(W)$ is a smooth convex function of $W$, and the $L_{2,1}$ penalty $|W|_{2,1}$ is not smooth. Like before, problem (3.9) is reformulated as a smooth constrained optimization problem

$$\arg\min_{(W,\mathbf{t}) \in C} \left\{ L(W) + \rho \sum_{i=1}^{G} t_i \right\} \tag{3.10}$$

where $C$ is the set $C = \{(W, \mathbf{t}) \mid \mathbf{w}_i \geq \mathbf{0}, |\mathbf{w}_i|_2 \leq t_i,\ i \in 1, 2, ..., G\}$, and $G$ is the number of groups in the group-sparsity regularizer. $C$ is a convex set, since it is the Cartesian product of intersections of convex sets (in particular, an ice-cream cone and the non-negative orthant), which must be convex.

The projection onto the convex set $C$ is given by the optimization problem

$$\pi_C(W, \mathbf{t}) = \arg\min_{(U,\mathbf{v}) \in C} \left\{ \frac{1}{2} |W - U|_F^2 + \frac{1}{2} |\mathbf{t} - \mathbf{v}|_2^2 \right\}. \tag{3.11}$$

As before, the objective in (3.11) is separated by group, yielding

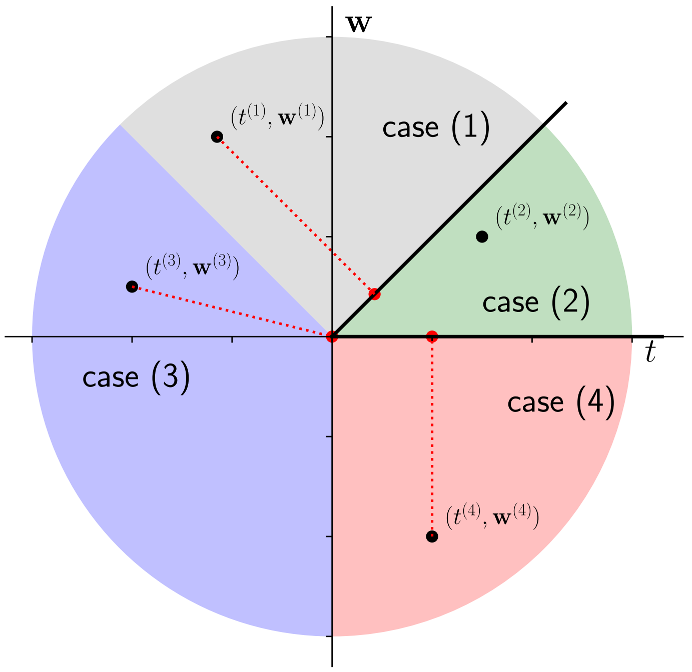


**Figure S2:** Projection onto the 1D non-negative subset of the ice-cream cone. There are now four cases: (1) $w_i \geq |t_i|$, corresponding to the gray shaded region; (2) $w_i \geq 0, w_i \leq t_i$, corresponding to the green shaded region; (3) $t_i < 0, w_i < -t_i$, corresponding to the blue shaded region; and (4) $w_i < 0, t_i > 0$, corresponding to the red shaded region. In cases (1), (3), and (4), $(w_i, t_i)$ lies outside the feasible region, and the associated red dotted lines and points correspond to the minimum norm projections onto the feasible set.

$$\pi_C(W, \mathbf{t}) = \arg \min_{(U,\mathbf{v}) \in C} \left\{ \sum_{i=1}^{G} \left[ \frac{1}{2} |\mathbf{w}_i - \mathbf{u}_i|^2 + \frac{1}{2}(t_i - v_i)^2 \right] \right\}. \tag{3.12}$$

Assuming that no variables are shared between groups, problem (3.12) can be solved separately for each group, yielding

$$\pi_{C_i}(\mathbf{w}_i, t_i) = \arg \min_{(\mathbf{u}_i, v_i) \in C} \left\{ \frac{1}{2} |\mathbf{w}_i - \mathbf{u}_i|^2 + \frac{1}{2}(t_i - v_i)^2 \right\}. \tag{3.13}$$

Consider the case where $\mathbf{w}_i$ is one-dimensional. A drawing of the two-dimensional $(\mathbf{w}_i, t_i)$ coordinate space is provided in Figure S2 to illustrate the four possible cases for projection:

1. $w_i \geq |t_i|$, corresponding to the gray shaded region of Figure S2. In this case, the solution is to orthogonally project $(w_i, t_i)$ onto the $w_i = v_i$ line.

2. $w_i \geq 0, w_i \leq t_i$, corresponding to the green shaded region of Figure S2. In this case, the point $(w_i, t_i)$ is already feasible, and so nothing needs to be done.

3. $t_i < 0, w_i < -t_i$, corresponding to the blue shaded region of Figure S2. In this case, the projection is performed by mapping $(w_i, t_i)$ onto the origin.

4. $w_i < 0, t_i > 0$, corresponding to the red shaded region of Figure S2. In this case, the projection is performed by mapping $(w_i, t_i)$ to $(0, t_i)$.

In mathematical terms, this corresponds to

$$u_i^* = \begin{cases} \frac{w_i+t_i}{2}, \ w_i \geq |t_i| & \text{(case 1)} \\ w_i, \ w_i \geq 0, w_i \leq t_i & \text{(case 2)} \\ 0, \ t_i < 0, w_i < -t_i & \text{(case 3)} \\ 0, \ w_i < 0, t_i > 0 & \text{(case 4)} \end{cases} \tag{3.14}$$

and

$$v_i^* = \begin{cases} \frac{w_i+t_i}{2}, \ w_i \geq |t_i| & \text{(case 1)} \\ t_i, \ w_i \geq 0, w_i \leq t_i & \text{(case 2)} \\ 0, \ t_i < 0, w_i < -t_i & \text{(case 3)} \\ t_i, \ w_i < 0, t_i > 0 & \text{(case 4).} \end{cases} \tag{3.15}$$

Unlike the unconstrained $L_{2,1}$ case, the projection is no longer rotationally symmetric about the $t_i$-axis because of the non-negativity constraint. Thus, generalizing the above to the $N$-dimensional case requires accounting for the possibility that some but not all of the components of $\mathbf{w}_i$ lie outside the non-negative quadrant.

The situation where some but not all of the components of $\mathbf{w}_i$ lie outside the non-negative quadrant is now examined in greater detail. Define $J_i^{(+)} = \{j \in 1, 2, \ldots, N \mid w_i^{(j)} \geq 0\}$ (e.g. the set of indices $j$ where $w_i^{(j)} \geq 0$), and define $J_i^{(-)} = \{j \in 1, 2, ..., N \mid w_i^{(j)} < 0\}$ (e.g. the set of indices $j$ where $w_i^{(j)} < 0$). All indices must belong to exactly one of $J_i^{(+)}$ or $J_i^{(-)}$. The single-group objective function from (3.13) is then split into

$$\frac{1}{2}|\mathbf{w}_i - \mathbf{u}_i|^2 + \frac{1}{2}(t_i - v_i)^2 = \frac{1}{2}\sum_{j \in J_i^{(+)}} [\mathbf{w}_i^{(j)} - \mathbf{u}_i^{(j)}]^2 + \frac{1}{2}\sum_{j \in J_i^{(-)}} [\mathbf{w}_i^{(j)} - \mathbf{u}_i^{(j)}]^2 + \frac{1}{2}(t_i - v_i). \tag{3.16}$$

First, consider only the variables $\{\mathbf{u}_i^{(j)} \mid j \in J_i^{(-)}\}$ (corresponding to the negative-valued components of $\mathbf{w}_i$ only). These variables are included only in the first term of (3.16). No matter the value of $t_i$, the optimal value for each of these variables $\{(\mathbf{u}_i^{(j)})^* \mid j \in J_i^{(-)}\}$ is exactly 0, corresponding to the lower $w < 0$ half of Figure S2. Intuitively, this is because all nonzero feasible values of $\{\mathbf{u}_i^{(j)} \mid j \in J_i^{(-)}\}$ are positive, and thus would increase the value of $[\mathbf{w}_i^{(j)} - \mathbf{u}_i^{(j)}]^2$.

Then, consider the remaining variables in the problem, $v_i$ and $\{\mathbf{u}_i^{(j)} \mid j \in J_i^{(+)}\}$. These variables lie in the upper $w \geq 0$ half of Figure S2, and thus, for these particular variables, the projection operator $\pi_D(\mathbf{w}, t)$ previously developed in Section 3.1.1 for the unconstrained regularized problem can be applied to solve for $v_i$ and $\{\mathbf{u}_i^{(j)} \mid j \in J_i^{(+)}\}$.

Letting max denote the element-wise maximum operator, the projection operator for the non-negative regularized problem can be expressed concisely as

$$\pi_C(\mathbf{w}_i, t_i) = \pi_D(\max(\mathbf{w}_i, \mathbf{0}), t_i). \tag{3.17}$$

The overall FISTA algorithm solving the non-negative $L_{2,1}$-regularized problem is given in Algorithm 4. Like before, $f_{\gamma, W_i, \mathbf{t}_i}(W_{i+1}, \mathbf{t}_{i+1})$ in the backtracking line search is defined as

$$f_{\gamma, W_i, \mathbf{t}_i}(W, \mathbf{t}) \triangleq f(W_i, \mathbf{t}_i) + \langle \boldsymbol{\nabla}'_W f(W', \mathbf{t}')|_{W'=W_i, \mathbf{t}'=\mathbf{t}_i}, W - W_i\rangle + \\ \langle \boldsymbol{\nabla}_{\mathbf{t}'} f(W', \mathbf{t}')|_{W'=W_i, \mathbf{t}'=\mathbf{t}_i}, \mathbf{t} - \mathbf{t}_i\rangle + \frac{\gamma}{2}|W - W_i|_F^2 + \frac{\gamma}{2}|\mathbf{t} - \mathbf{t}_i|_2^2.$$

**Algorithm 4** FISTA for non-negative $L_{2,1}$-regularized problem

Hyperparameter $\rho_{L21}$
**procedure** NonNegative-FISTA-L21($L(W)$, $W_0$, $G$, $\gamma$)
  Randomly initialize variable $\mathbf{t} \in \mathbb{R}^{|G|}_{\geq 0}$
  $f(W, \mathbf{t}) \triangleq L(W) + \rho_{L21} \mathbf{1}^T \mathbf{t}$
  Initialize $i = 1, W_1 = W_0, b_{-1} = 0, b_0 = 1$
  $W^\star = W_0$
  **repeat**
    $\alpha_i = \frac{b_{i-2} - 1}{b_{i-1}}$
    $W_i' = W_i + \alpha_i (W_i - W_{i-1})$
    $\mathbf{t}_i' = \mathbf{t}_i + \alpha_i (\mathbf{t}_i - \mathbf{t}_{i-1})$
    **while** True **do** ▷ Backtracking line search
      $\gamma = \frac{\gamma}{2}$
      $\bar{W}_{i+1} = W_i' - \frac{1}{\gamma} \boldsymbol{\nabla}_W f(W, \mathbf{t})\big|_{W = W_i', \mathbf{t} = \mathbf{t}_i'}$
      $\bar{\mathbf{t}}_{i+1} = \mathbf{t}_i' - \frac{1}{\gamma} \boldsymbol{\nabla}_{\mathbf{t}} f(W, \mathbf{t})\big|_{W = W_i', \mathbf{t} = \mathbf{t}_i'}$
      $W_{i+1}, \mathbf{t}_{i+1} = \pi_{D_G}(\max(\bar{W}_{i+1}, 0), \bar{\mathbf{t}}_{i+1})$
      **if** $f(W_{i+1}, \mathbf{t}_{i+1}) \leq f_{\gamma, W_i, \mathbf{t}_i}(W_{i+1}, \mathbf{t}_{i+1})$ **then**
        $W^\star = W_{i+1}$
        break
      **end if**
    **end while**
    $b_i = \frac{1 + \sqrt{1 + 4 b_{i-1}^2}}{2}$
  **until** Converged
  **return** $W^\star$
**end procedure**

#### Fourier Waveform Optimization

#### 4.1 Solving waveform shapes linear system in Fourier domain

This section describes the construction of the linear system of equations associated with the first-order conditions of optimality for the waveform shapes fitting step. This optimization is performed in the Fourier domain, to simplify accounting for the basis waveform temporal shifts, and thus the coefficients of the linear system are found by differentiating the mean square error loss function with respect to the real and imaginary components of the basis waveforms.

The construction of the linear system is rather tedious; for simplicity we start by first finding the linear coefficients for the unregularized waveform shapes problem (without the Gaussian shape prior), and then derive the modifications needed to solve the regularized waveform shapes problem (with the Gaussian shape prior).

##### 4.1.1 Solving for basis waveforms without the Gaussian shape prior

Let $\tilde{B}$ denote the discrete Fourier transform (DFT) of the basis waveforms $B$, $\tilde{X}_{:,l}$ the DFT of the $l^{\text{th}}$ observed data waveform, $\tilde{P}_l$ the complex-valued Fourier phase shift matrix corresponding to the previously-estimated time shifts for the $l^{\text{th}}$ data waveform, $A_{:,l}$ the previously-estimated amplitudes for the $l^{\text{th}}$ data waveform, and $\odot$ Hadamard element-wise matrix multiplication. The least squares optimization problem in frequency domain over all observed data waveforms for a given cell is

$$\arg\min_{\tilde{B}} \left\{ \sum_{l=1}^{L} \frac{1}{2} |(\tilde{B} \odot \tilde{P^{(l)}}) A_{:,l} - \tilde{X}_{:,l}|_2^2 \right\}. \tag{4.1}$$

This is an unconstrained convex minimization problem, and has first order condition of optimality

$$\nabla_{\tilde{B}} \left\{ \sum_{l=1}^{L} \frac{1}{2} |(\tilde{B} \odot \tilde{P^{(l)}} A_{:,l} - \tilde{X}_{:,l}|_2^2 \right\} = 0. \tag{4.2}$$

Interchanging summation and the gradient operator leaves

$$\sum_{l=1}^{L} \nabla_{\tilde{B}} \left\{ \frac{1}{2} |(\tilde{B} \odot \tilde{P^{(l)}}) A_{:,l} - \tilde{X}_{:,l}|_2^2 \right\} = 0. \tag{4.3}$$

Let the subscript Re correspond to the real component of a complex number (e.g. $\tilde{y}_{\text{Re}} = \text{Re}\{\tilde{y}\}$), and let the subscript Im correspond to the imaginary component of a complex number (e.g. $\tilde{y}_{\text{Im}} = \text{Im}\{\tilde{y}\}$). Furthermore, define $\mathscr{L}_l \triangleq \frac{1}{2}|(\tilde{B} \odot \tilde{P^{(l)}}) A_{:,l} - \tilde{X}_{:,l}|_2^2$ as the contribution of data waveform $l$ to the overall mean square error. Taking the derivative of $\mathscr{L}_l$ with respect to $(\tilde{B}_{\text{Re}})_{f,k}$, the real component of the $f^{\text{th}}$ Fourier coefficient of the $k^{\text{th}}$ basis waveform, results in

$$\begin{aligned} \frac{\partial \mathscr{L}_l}{\partial (\tilde{B}_{\text{Re}})_{f,k}} = A_{k,l} \sum_j & \left\{ (\tilde{B}_{\text{Re}})_{f,j} A_{j,l} [(\tilde{P}^{(l)}_{\text{Re}})_{f,k} (\tilde{P}^{(l)}_{\text{Re}})_{f,j} + (\tilde{P}^{(l)}_{\text{Im}})_{f,k} (\tilde{P}^{(l)}_{\text{Im}})_{f,j}] \right\} \\ + A_{k,l} \sum_j & \left\{ (\tilde{B}_{\text{Im}})_{f,j} A_{j,l} [-(\tilde{P}^{(l)}_{\text{Re}})_{f,k} (\tilde{P}^{(l)}_{\text{Im}})_{f,j} + (\tilde{P}^{(l)}_{\text{Im}})_{f,k} (\tilde{P}^{(l)}_{\text{Re}})_{f,j}] \right\} \\ & - (\tilde{P}^{(l)}_{\text{Re}})_{f,k} (\tilde{X}_{\text{Re}})_{f,l} A_{k,l} - (\tilde{P}^{(l)}_{\text{Im}})_{f,k} (\tilde{X}_{\text{Im}})_{f,l} A_{k,l}. \end{aligned} \tag{4.4}$$

Similarly, taking the derivative of $\mathscr{L}_l$ with respect to $(\tilde{B}_{\text{Im}})_{f,k}$, the imaginary component of the $f^{\text{th}}$ Fourier coefficient of the $k^{\text{th}}$ basis waveform, results in

$$\frac{\partial \mathscr{L}_l}{\partial (\tilde{B}_{\text{Im}})_{f,k}} = A_{k,l} \sum_j \left\{ (\tilde{B}_{\text{Re}})_{f,j} A_{j,l} [(\tilde{P}^{(l)}_{\text{Re}})_{f,k} (\tilde{P}^{(l)}_{\text{Im}})_{f,j} - (\tilde{P}^{(l)}_{\text{Im}})_{f,k} (\tilde{P}^{(l)}_{\text{Re}})_{f,j}] \right\}$$
$$+ A_{k,l} \sum_j \left\{ (\tilde{B}_{\text{Im}})_{f,j} A_{j,l} [(\tilde{P}^{(l)}_{\text{Im}})_{f,k} (\tilde{P}^{(l)}_{\text{Im}})_{f,j} + (\tilde{P}^{(l)}_{\text{Re}})_{f,k} (\tilde{P}^{(l)}_{\text{Re}})_{f,j}] \right\}$$
$$- (\tilde{P}^{(l)}_{\text{Re}})_{f,k} (\tilde{X}_{\text{Im}})_{f,l} A_{k,l} + (\tilde{P}^{(l)}_{\text{Im}})_{f,k} (\tilde{X}_{\text{Re}})_{f,l} A_{k,l}. \tag{4.5}$$

**Constructing linear equations from the first-order conditions**

The optimal waveform shapes are found by solving a system of linear equations in terms of $(\tilde{B}_{\text{Re}})_{f,k}$ and $(\tilde{B}_{\text{Im}})_{f,k}$ corresponding to the first-order conditions of optimality. Two families of equations are constructed, the first by differentiating with respect to the $\tilde{B}_{\text{Re}}$, and the second by differentiating with respect to $\tilde{B}_{\text{Im}}$.

**First equation family: differentiating with respect to $\tilde{B}_{\text{Re}}$** Consider the first-order condition associated with differentiating the objective with respect to $(\tilde{B}_{\text{Re}})_{f,k}$, the real component of the $f^{\text{th}}$ Fourier coefficient of the $k^{\text{th}}$ basis waveform:

$$\sum_{l=1}^{L} \frac{\partial}{\partial (\tilde{B}_{\text{Re}})_{f,k}} \left\{ \frac{1}{2} |(\tilde{B} \odot \tilde{P^{(l)}}) A_{:,l} - \tilde{X}_{:,l}|_2^2 \right\} = \sum_{l=1}^{L} \frac{\partial \mathscr{L}_l}{\partial (\tilde{B}_{\text{Re}})_{f,k}} = 0. \tag{4.6}$$

$(\tilde{B}_{\text{Re}})_{f,k}$ and $(\tilde{B}_{\text{Im}})_{f,k}$ for each of the basis waveforms $k \in \{1, 2, ..., K\}$ appear in this equation, and hence there are $2K$ unknowns in this equation. Let $[(\tilde{B}_{\text{Re}})_{f,k}]^{(\text{Re},f,k)}$ denote the coefficient of $(\tilde{B}_{\text{Re}})_{f,k}$, and $[(\tilde{B}_{\text{Im}})_{f,k}]^{(\text{Re},f,k)}$ denote the coefficient of $(\tilde{B}_{\text{Im}})_{f,k}$ in this equation, where the superscript $(\text{Re}, f, k)$ serves as a reminder that the equation was constructed by differentiating with respect to the real component of the $f^{\text{th}}$ Fourier coefficient of the $k^{\text{th}}$ basis waveform. The coefficients of the equation are

$$[(\tilde{B}_{\text{Re}})_{f,k}]^{(\text{Re},f,k)} = \sum_{l=1}^{L} \left\{ A_{k,l} A_{j,l} [(\tilde{P}^{(l)}_{\text{Re}})_{f,k} (\tilde{P}^{(l)}_{\text{Re}})_{f,j} + (\tilde{P}^{(l)}_{\text{Im}})_{f,k} (\tilde{P}^{(l)}_{\text{Im}})_{f,j}] \right\} \tag{4.7}$$

and

$$[(\tilde{B}_{\text{Im}})_{f,k}]^{(\text{Re},f,k)} = \sum_{l=1}^{L} \left\{ A_{k,l} A_{j,l} [(\tilde{P}^{(l)}_{\text{Im}})_{f,k} (\tilde{P}^{(l)}_{\text{Re}})_{f,j} - (\tilde{P}^{(l)}_{\text{Re}})_{f,k} (\tilde{P}^{(l)}_{\text{Im}})_{f,j}] \right\} \tag{4.8}$$

The right hand side of the equation $R^{(\text{Re},f,k)}$ is constructed from the third terms of each of equations (4.4) and (4.5):

$$R^{(\text{Re},f,k)} = \sum_{l=1}^{L} \left\{ (\tilde{P}^{(l)}_{\text{Re}})_{f,k} (\tilde{X_{\text{Re}}})_{f,l} A_{k,l} + (\tilde{P}^{(l)}_{\text{Im}})_{f,k} (\tilde{X_{\text{Im}}})_{f,l} A_{k,l} \right\} \tag{4.9}$$

Thus, the overall linear equation associated with the derivative with respect to $(\tilde{B}_{\text{Re}})_{f,k}$ is

$$\sum_{j=1}^{K} [(\tilde{B}_{\text{Re}})_{f,j}]^{(\text{Re},f,k)} (\tilde{B}_{\text{Re}})_{f,j} + \sum_{j=1}^{K} [(\tilde{B}_{\text{Im}})_{f,j}]^{(\text{Re},f,k)} (\tilde{B}_{\text{Im}})_{f,j} = R^{(\text{Re},f,k)}. \tag{4.10}$$

**Second equation family: differentiating with respect to $\tilde{B}_{\text{Im}}$** Consider the first-order condition associated with differentiating the objective with respect to $(\tilde{B}_{\text{Im}})_{f,k}$, the imaginary component of the $f^{\text{th}}$ Fourier coefficient of the $k^{\text{th}}$ basis waveform:

$$\sum_{l=1}^{L} \frac{\partial}{\partial (\tilde{B}_{\text{Im}})_{f,k}} \left\{ \frac{1}{2} |(\tilde{B} \odot \tilde{P^{(l)}}) A_{:,l} - \tilde{X}_{:,l}|_2^2 \right\} = \sum_{l=1}^{L} \frac{\partial \mathscr{L}_l}{\partial (\tilde{B}_{\text{Im}})_{f,k}} = 0. \tag{4.11}$$

$(\tilde{B}_{\text{Re}})_{f,k}$ and $(\tilde{B}_{\text{Im}})_{f,k}$ for each of the basis waveforms $k \in \{1, 2, ..., K\}$ appear in this equation, and hence there are $2K$ unknowns in this equation. Let $[(\tilde{B}_{\text{Re}})_{f,k}]^{(\text{Im},f,k)}$ denote the coefficient of $(\tilde{B}_{\text{Re}})_{f,k}$, and $[(\tilde{B}_{\text{Im}})_{f,k}]^{(\text{Im},f,k)}$ denote the coefficient of $(\tilde{B}_{\text{Im}})_{f,k}$ in this equation, where the superscript $(\text{Im}, f, k)$ serves as a reminder that the equation was constructed by differentiating with respect to the imaginary component of the $f^{\text{th}}$ Fourier coefficient of the $k^{\text{th}}$ basis waveform. The coefficients of the equation are

$$[(\tilde{B}_{\text{Re}})_{f,k}]^{(\text{Im},f,k)} = \sum_{l=1}^{L} \left\{ A_{k,l} A_{j,l} [(\tilde{P}_{\text{Re}}^{(l)})_{f,k} (\tilde{P}_{\text{Im}}^{(l)})_{f,j} - (\tilde{P}_{\text{Im}}^{(l)})_{f,k} (\tilde{P}_{\text{Re}}^{(l)})_{f,j}] \right\} \tag{4.12}$$

and

$$[(\tilde{B}_{\text{Im}})_{f,k}]^{(\text{Im},f,k)} = \sum_{l=1}^{L} \{ A_{k,l} A_{j,l} [(\tilde{P}_{\text{Im}}^{(l)})_{f,k} (\tilde{P}_{\text{Im}}^{(l)})_{f,j} + (\tilde{P}_{\text{Re}}^{(l)})_{f,k} (\tilde{P}_{\text{Re}}^{(l)})_{f,j}] \} \tag{4.13}$$

The right hand side of the equation $R^{(\text{Im},f,k)}$ is constructed from the third terms of each of equations (4.4) and (4.5):

$$R^{(\text{Im},f,k)} = \sum_{l} \left\{ (\tilde{P}_{\text{Re}}^{(l)})_{f,k} (\tilde{X}_{\text{Im}})_{f,l} A_{k,l} - (\tilde{P}_{\text{Im}}^{(l)})_{f,k} (\tilde{X}_{\text{Re}})_{f,l} A_{k,l} \right\} \tag{4.14}$$

Thus, the overall linear equation associated with the derivative with respect to $(\tilde{B}_{\text{Im}})_{f,k}$ is

$$\sum_{j=1}^{K} [(\tilde{B}_{\text{Re}})_{f,j}]^{(\text{Im},f,k)} (\tilde{B}_{\text{Re}})_{f,j} + \sum_{j=1}^{K} [(\tilde{B}_{\text{Im}})_{f,j}]^{(\text{Im},f,k)} (\tilde{B}_{\text{Im}})_{f,j} = R^{(\text{Im},f,k)}. \tag{4.15}$$

**Constructing $F$ systems of $2K \times 2K$ equations** Differentiating with respect to the real and imaginary components of each of the $F$ Fourier coefficients for each of the $K$ basis waveforms produces $F \times 2K$ linear equations in $F \times 2K$ variables. Careful inspection of equations (4.10) and (4.15) shows that a separate system of $2K \times 2K$ linear equations can be constructed for each frequency $f$ (in other words, the overall linear system of equations is block-diagonal). Thus, the basis waveforms $\tilde{B}$ can be computed by solving $F$ distinct linear systems, each with $2K$ variables and $2K$ equations.

##### 4.1.2 Solving for basis waveforms with the Gaussian shape prior

The linear system of equations for determining basis waveform shapes described in Section 4.1.2 becomes under-determined or extremely ill-conditioned if at least one of basis waveforms is not observed for a particular cell (occurring, for instance, when the recorded cell lies near the edge of the electrode array). A Gaussian waveform shape prior that weakly constrains the learned waveform shapes to approximately resemble known waveform shapes is introduced to address this problem.

Let column vectors $\mathbf{b}_1, \mathbf{b}_2, ..., \mathbf{b}_K \in \mathbb{R}^T$ denote each of the compartment basis waveforms in time domain. The Gaussian prior for each of the basis waveforms has form

$$\mathbf{b}_k \sim \mathcal{N}(\mu_k, \Sigma_k) \tag{4.16}$$

where $\mu_i$ is the prior mean waveform of the specific compartment (e.g. a typical somatic waveform), and $\Sigma$ is a temporal covariance matrix. The entries of temporal covariance matrix can be specified using a 1D Gaussian process kernel, for example, an exponentiated quadratic kernel $k(t_1, t_2) = \sigma^2 \exp\{-|t_1 - t_2|^2/(2l^2)\}$.

Like in Section 4.1.2, the waveform shape optimization with the Gaussian prior is performed in the Fourier domain, and thus the prior regularization penalty must be translated to the Fourier domain. Define $F$ as the DFT analysis matrix for computing the one-dimensional Fourier transform of real-valued input. $F$ is a complex-valued $(\lfloor \frac{T}{2} \rfloor + 1) \times T$ matrix, where $T$ is the number of samples in time domain for the basis waveforms. We further define $F_{\text{Re}} \triangleq \text{Re}\{F\}$ and $F_{\text{Im}} \triangleq \text{Im}\{F\}$ as the real and imaginary components of the matrix $F$, respectively. $F_{\text{Re}}$ is a real-valued $\lfloor \frac{T}{2} \rfloor + 1) \times T$ matrix, and $F_{\text{Im}}$ is a real-valued $(\lfloor \frac{T-1}{2} \rfloor \times T$ matrix. Note that splitting $F$ into real and imaginary components is only useful in the present case when the signal $\mathbf{b}$ is real-valued, since otherwise separating the real and imaginary components of the Fourier transform would not be linear operations.

Because the basis waveforms $\mathbf{b}$ are real-valued, the real component of the basis waveform Fourier transform is $\tilde{\mathbf{b}}_{\text{Re}} = F_{\text{Re}}\mathbf{b}$, and the imaginary component is $\tilde{\mathbf{b}}_{\text{Im}} = F_{\text{Im}}\mathbf{b}$. Define the real-valued vector $\mathbf{z} \triangleq [\tilde{\mathbf{b}}_{\text{Re}}^T\ \tilde{\mathbf{b}}_{\text{Im}}^T]^T$, and the $T \times T$ real-valued matrix $G$

$$G \triangleq \begin{bmatrix} F_{\text{Re}} \\ F_{\text{Im}} \end{bmatrix} \tag{4.17}$$

Because $\mathbf{z} = G\mathbf{b}$ is a linear transformation of the Gaussian random variable $\mathbf{b}$, $\mathbf{z}$ is also a jointly Gaussian random variable, and is distributed according to

$$\mathbf{z} \sim \mathscr{N}(G\mu, G\Sigma G^T) \tag{4.18}$$

which has pdf

$$p(\mathbf{z}) \propto \exp\{-\frac{1}{2}(\mathbf{z} - G\mu)^T(G_{\text{stack}}\Sigma G^T)^{-1}(\mathbf{z} - G\mu)\}. \tag{4.19}$$

The Gaussian prior penalty term associated basis waveform $j$, $\mathscr{P}_j$, is then rewritten as

$$\mathscr{P}_j \triangleq \frac{\lambda_p}{2}(\mathbf{z}_j - G\mu_j)^T(G\Sigma_k G^T)^{-1}(\mathbf{z}_j - G\mu_j) \tag{4.20}$$

and therefore the overall waveform shapes objective function, consisting of the sum of the mean square error terms and the Gaussian prior penalty terms, is

$$\sum_{l=1}^{L} \frac{1}{2}|(\tilde{B} \odot \tilde{P}_l)A_{:,l} - \tilde{X}_{:,l}|_2^2 + \frac{\lambda_p}{2}\sum_{j=1}^{K} \mathscr{P}_j. \tag{4.21}$$

**Constructing linear equations from the first-order conditions**

The linear system of equations from Section 4.1.2 must be updated to include the Gaussian prior penalty. Specifically, the contributions from the gradient of Gaussian prior penalty must be incorporated into the first-order conditions of optimality. The gradient of the Gaussian prior penalty with respect to the basis waveform Fourier coefficients $\mathbf{z}_k$ is

$$\boldsymbol{\nabla}_{\mathbf{z}_k}\left\{\frac{\lambda_p}{2}\sum_{j=1}^{K} \mathscr{P}_j\right\} = \lambda_p(G\Sigma_k G^T)^{-1}(\mathbf{z}_k - G\mu_k). \tag{4.22}$$

A notational difference regarding the indexing of the basis waveform Fourier coefficients must be resolved in order to incorporate the gradients in equation (4.22) with the first-order condition equations (4.10) and (4.10). Specifically, $(\tilde{B}_{\text{Re}})_{f,k} = \{\mathbf{z}_k\}_{(\text{Re},f)}$ and $(\tilde{B}_{\text{Im}})_{f,k} = \{\mathbf{z}_k\}_{(\text{Im},f)}$, where the subscripts $(\text{Re}, f)$ and $(\text{Im}, f)$ represent the indices of the entries in $\mathbf{z}_k$ corresponding to the real and imaginary components of the Fourier coefficient at frequency $f$, respectively. Similarly, let $\{(G\Sigma_k G^T)^{-1}\}_{(\text{Re},f),(\text{Im},g)}$ denote the entry at the $(\text{Re}, f)$ row and $(\text{Im}, g)$ column of the matrix $(G\Sigma_k G^T)^{-1}$, e.g. the entry at the $m^{\text{th}}$ row and $n^{\text{th}}$ column, where $m$ is the index of the entry in $\mathbf{z}_k$ corresponding to the real component of the Fourier coefficient at frequency $f$, and $n$ is the index of the entry in $\mathbf{z}_k$ corresponding to the imaginary component of the Fourier coefficient at frequency $g$.

**First equation family, differentiating with respect to $(\tilde{B}_{\text{Re}})_{f,k}$, a.k.a. $\{\mathbf{z}_k\}_{(\text{Re},f)}$** Differentiating the Gaussian prior penalty with respect to $(\tilde{B}_{\text{Re}})_{f,k}$, a.k.a. $(\mathbf{z}_k)_{(\text{Re},f)}$, corresponds to the $(\text{Re}, f)^{\text{th}}$ entry of (4.22):

$$
\begin{aligned}
\frac{\partial}{\partial(\tilde{B}_{\mathrm{Re}})_{f,k}}\left\{\frac{\lambda_p}{2}\sum_{j=1}^{K}\mathscr{P}_j\right\} = \lambda_p \sum_g &\{(G\Sigma_k G^T)^{-1}\}_{(\mathrm{Re},f),(\mathrm{Re},g)}(\tilde{B}_{\mathrm{Re}})_{g,k} \\
+\lambda_p \sum_g &\{(G\Sigma_k G^T)^{-1}\}_{(\mathrm{Re},f),(\mathrm{Im},g)}(\tilde{B}_{\mathrm{Im}})_{g,k} \\
-\lambda_p \sum_g &\{(G\Sigma_k G^T)^{-1}\}_{(\mathrm{Re},f),(\mathrm{Re},g)}\{G\mu_k\}_{(\mathrm{Re},f)} \\
-\lambda_p \sum_g &\{(G\Sigma_k G^T)^{-1}\}_{(\mathrm{Re},f),(\mathrm{Im},g)}\{G\mu_k\}_{(\mathrm{Im},g)}
\end{aligned}
\tag{4.23}
$$

The first and second summations in equation (4.26) contain the variables (the entries $\tilde{B}$), while the third and fourth summations in (4.26) are constants. Defining

$$
\begin{aligned}
R_{\mathrm{prior}}^{(\mathrm{Re},f,k)} \triangleq \lambda_p \sum_g &\{(G\Sigma_k G^T)^{-1}\}_{(\mathrm{Re},f),(\mathrm{Re},g)}\{G\mu_k\}_{(\mathrm{Re},f)} \\
+\lambda_p \sum_g &\{(G\Sigma_k G^T)^{-1}\}_{(\mathrm{Re},f),(\mathrm{Im},g)}\{G\mu_k\}_{(\mathrm{Im},g)},
\end{aligned}
\tag{4.24}
$$

inclusion of the Gaussian prior into equation (4.10) results in

$$
\begin{aligned}
\sum_{j=1}^{K}[(\tilde{B}_{\mathrm{Re}})_{f,j}]^{(\mathrm{Re},f,k)}(\tilde{B}_{\mathrm{Re}})_{f,j} + \sum_{j=1}^{K}[(\tilde{B}_{\mathrm{Im}})_{f,j}]^{(\mathrm{Re},f,k)}(\tilde{B}_{\mathrm{Im}})_{f,j}& \\
+\lambda_p \sum_g \{(G\Sigma_k G^T)^{-1}\}_{(\mathrm{Re},f),(\mathrm{Re},g)}(\tilde{B}_{\mathrm{Re}})_{g,k}& \\
+\lambda_p \sum_g \{(G\Sigma_k G^T)^{-1}\}_{(\mathrm{Re},f),(\mathrm{Im},g)}(\tilde{B}_{\mathrm{Im}})_{g,k} = R^{(\mathrm{Re},f,k)} + R_{\mathrm{prior}}^{(\mathrm{Re},f,k)}&.
\end{aligned}
\tag{4.25}
$$

**Second equation family, differentiating with respect to $(\tilde{B}_{\mathrm{Im}})_{f,k}$, a.k.a. $\{\mathbf{z}_k\}_{(\mathrm{Im},f)}$** Differentiating the Gaussian prior penalty with respect to $(\tilde{B}_{\mathrm{Im}})_{f,k}$, a.k.a. $\{\mathbf{z}_k\}_{(\mathrm{Im},f)}$ corresponds to the $(\mathrm{Im}, f)^{\mathrm{th}}$ entry of (4.22):

$$
\begin{aligned}
\frac{\partial}{\partial(\tilde{B}_{\mathrm{Im}})_{f,k}}\left\{\frac{\lambda_p}{2}\sum_{j=1}^{K}\mathscr{P}_j\right\} = \lambda_p \sum_g &\{(G\Sigma_k G^T)^{-1}\}_{(\mathrm{Im},f),(\mathrm{Re},g)}(\tilde{B}_{\mathrm{Re}})_{g,k} \\
+\lambda_p \sum_g &\{(G\Sigma_k G^T)^{-1}\}_{(\mathrm{Im},f),(\mathrm{Im},g)}(\tilde{B}_{\mathrm{Im}})_{g,k} \\
-\lambda_p \sum_g &\{(G\Sigma_k G^T)^{-1}\}_{(\mathrm{Im},f),(\mathrm{Re},g)}\{G\mu_k\}_{(\mathrm{Re},f)} \\
-\lambda_p \sum_g &\{(G\Sigma_k G^T)^{-1}\}_{(\mathrm{Im},f),(\mathrm{Im},g)}\{G\mu_k\}_{(\mathrm{Im},g)}
\end{aligned}
\tag{4.26}
$$

Defining

$$
\begin{aligned}
R_{\mathrm{prior}}^{(\mathrm{Im},f,k)} \triangleq \lambda_p \sum_g &\{(G\Sigma_k G^T)^{-1}\}_{(\mathrm{Im},f),(\mathrm{Re},g)}\{G\mu_k\}_{(\mathrm{Re},f)} \\
+\lambda_p \sum_g &\{(G\Sigma_k G^T)^{-1}\}_{(\mathrm{Im},f),(\mathrm{Im},g)}\{G\mu_k\}_{(\mathrm{Im},g)},
\end{aligned}
\tag{4.27}
$$

and thus inclusion of the Gaussian prior into (4.15) results in

$$
\begin{aligned}
\sum_{j=1}^{K}[(\tilde{B}_{\mathrm{Re}})_{f,j}]^{(\mathrm{Im},f,k)}(\tilde{B}_{\mathrm{Re}})_{f,j} + \sum_{j=1}^{K}[(\tilde{B}_{\mathrm{Im}})_{f,j}]^{(\mathrm{Im},f,k)}(\tilde{B}_{\mathrm{Im}})_{f,j} \\
+\lambda_p \sum_{g}\{(G\Sigma_k G^T)^{-1}\}_{(\mathrm{Im},f),(\mathrm{Re},g)}(\tilde{B}_{\mathrm{Re}})_{g,k} \\
+\lambda_p \sum_{g}\{(G\Sigma_k G^T)^{-1}\}_{(\mathrm{Im},f),(\mathrm{Im},g)}(\tilde{B}_{\mathrm{Im}})_{g,k} = R^{(\mathrm{Im},f,k)} + R^{(\mathrm{Im},f,k)}_{\mathrm{prior}}.
\end{aligned}
\tag{4.28}
$$

**Constructing a single $KN \times KN$ system of equations** The Gaussian prior penalty couples coefficients of all frequencies for the same basis waveform together, and thus the block diagonal structure from Section is lost. Solution of the first-order conditions with the Gaussian prior penalty therefore requires solving a single $KN \times KN$ system of linear equations.
